# Highly-accurate long-read sequencing improves variant detection and assembly of a human genome

**DOI:** 10.1101/519025

**Authors:** Aaron M. Wenger, Paul Peluso, William J. Rowell, Pi-Chuan Chang, Richard J. Hall, Gregory T. Concepcion, Jana Ebler, Arkarachai Fungtammasan, Alexey Kolesnikov, Nathan D. Olson, Armin Töpfer, Michael Alonge, Medhat Mahmoud, Yufeng Qian, Chen-Shan Chin, Adam M. Phillippy, Michael C. Schatz, Gene Myers, Mark A. DePristo, Jue Ruan, Tobias Marschall, Fritz J. Sedlazeck, Justin M. Zook, Heng Li, Sergey Koren, Andrew Carroll, David R. Rank, Michael W. Hunkapiller

## Abstract

The major DNA sequencing technologies in use today produce either highly-accurate short reads or noisy long reads. We developed a protocol based on single-molecule, circular consensus sequencing (CCS) to generate highly-accurate (99.8%) long reads averaging 13.5 kb and applied it to sequence the well-characterized human HG002/NA24385. We optimized existing tools to comprehensively detect variants, achieving precision and recall above 99.91% for SNVs, 95.98% for indels, and 95.99% for structural variants. We estimate that 2,434 discordances are correctable mistakes in the high-quality Genome in a Bottle benchmark. Nearly all (99.64%) variants are phased into haplotypes, which further improves variant detection. *De novo* assembly produces a highly contiguous and accurate genome with contig N50 above 15 Mb and concordance of 99.998%. CCS reads match short reads for small variant detection, while enabling structural variant detection and *de novo* assembly at similar contiguity and markedly higher concordance than noisy long reads.

## Introduction

DNA sequencing technologies have improved at rates eclipsing Moore’s law^1^ revolutionizing biological sciences. Beginning in the 1970s, Sanger sequencing^2^, and subsequent automation^3^ facilitated large scale DNA sequencing projects and paved the way for modern genomic research^4–7^. The first reference genomes were followed by the advent of several high-throughput sequencing technologies (next-generation sequencing or NGS) including 454™, Solexa/Illumina^®^, ABI^®^ Solid™, Complete Genomics™, and Ion Torrent™. These technologies employed a range of chemistries and detection strategies^8–13^. All produce relatively accurate reads but are limited in read length, typically to less than 300 basepairs (bp). These accurate short reads are well-suited for calling single-nucleotide variants (SNVs) and small insertions and deletions (indels), but are lacking for long-range applications such as *de novo* assembly, haplotype phasing, and structural variant detection.

For these applications, vastly superior results^14–18^ are obtained with technologies like PacBio^®^ SMRT Sequencing^19^ and Oxford Nanopore sequencing^20^ that produce long reads (~10 kb). These technologies rely on single-molecule detection and are characterized by reduced read accuracy (75-90%)^19,20^. High consensus accuracy has been demonstrated through read-to-read error correction, but the process is computationally intensive, and errors remain from mis-mapping reads and mixing haplotypes during correction^15,21^. As a result of the error rate, long-read technologies are rarely used to detect SNVs and indels.

Today, human genomes are sequenced at population scales, but it remains necessary to combine sequencing technologies to cover all types of genetic variation, which increases cost and adds complexity to projects. A sequencing technology with long read length and high accuracy would enable a single experiment for comprehensive variant discovery.

Recent gains in read length for SMRT Sequencing and optimized DNA template preparation suggested an opportunity to unify high accuracy with long read lengths using circular consensus sequencing (CCS)^22,23^. CCS derives a consensus sequence from multiple passes of a single template molecule, producing accurate reads from noisy individual subreads.

Here, we highlight the performance of these highly-accurate, long CCS™ reads by sequencing and analyzing the well-characterized human male HG002/NA24385^24,25^. The HG002 sample is one of the benchmark samples available from the Genome in a Bottle (GIAB) Consortium. GIAB provides physical reference materials along with detailed characterization of the sample genome, defining “high-confidence regions” at which the sequence of the sample is known and “high-confidence variants” within those regions at which the sample differs from the human reference genome. Thus, it is an ideal sample for study of sequencing accuracy and variant detection. We apply and extend standard analysis tools to identify variation in HG002, demonstrating performance that rivals or surpasses existing technologies for small and large variation detection as well as genome assembly and haplotype phasing.

## Results

### CCS Library Preparation and Sequencing

A SMRTbell library tightly distributed at 15 kb was chosen for circular consensus sequencing (**Figure 1a**, **Supplementary Figure 1**) based on estimates of 150 kb polymerase read length and a requirement of 10 passes to achieve Q30 read accuracy (**Figure 1b**). CCS reads with a predicted accuracy of at least Q20 (99%) were retained (**Supplementary Figure 2a**). The total CCS read yield was 89 Gb (2.3±0.4 Gb over 39 SMRT Cells), with read length of 13.5±1.2 kb (**Figure 1c**). The predicted accuracy of the CCS reads has a median of Q30 (99.9%) and a mean of Q27 (99.8%) (**Figure 1c**). Predicted accuracy matches well with concordance to the GIAB HG002 benchmark (average [Q_predicted_ – Q_concordance_] = −1.2), which indicates that the predicted accuracy is well calibrated (**Supplementary Figure 2b-c**). Average mapped coverage of the genome is 28-fold, with minimal difference across [GC] content (**Supplementary Figure 2d-e**).

**Figure 1.**
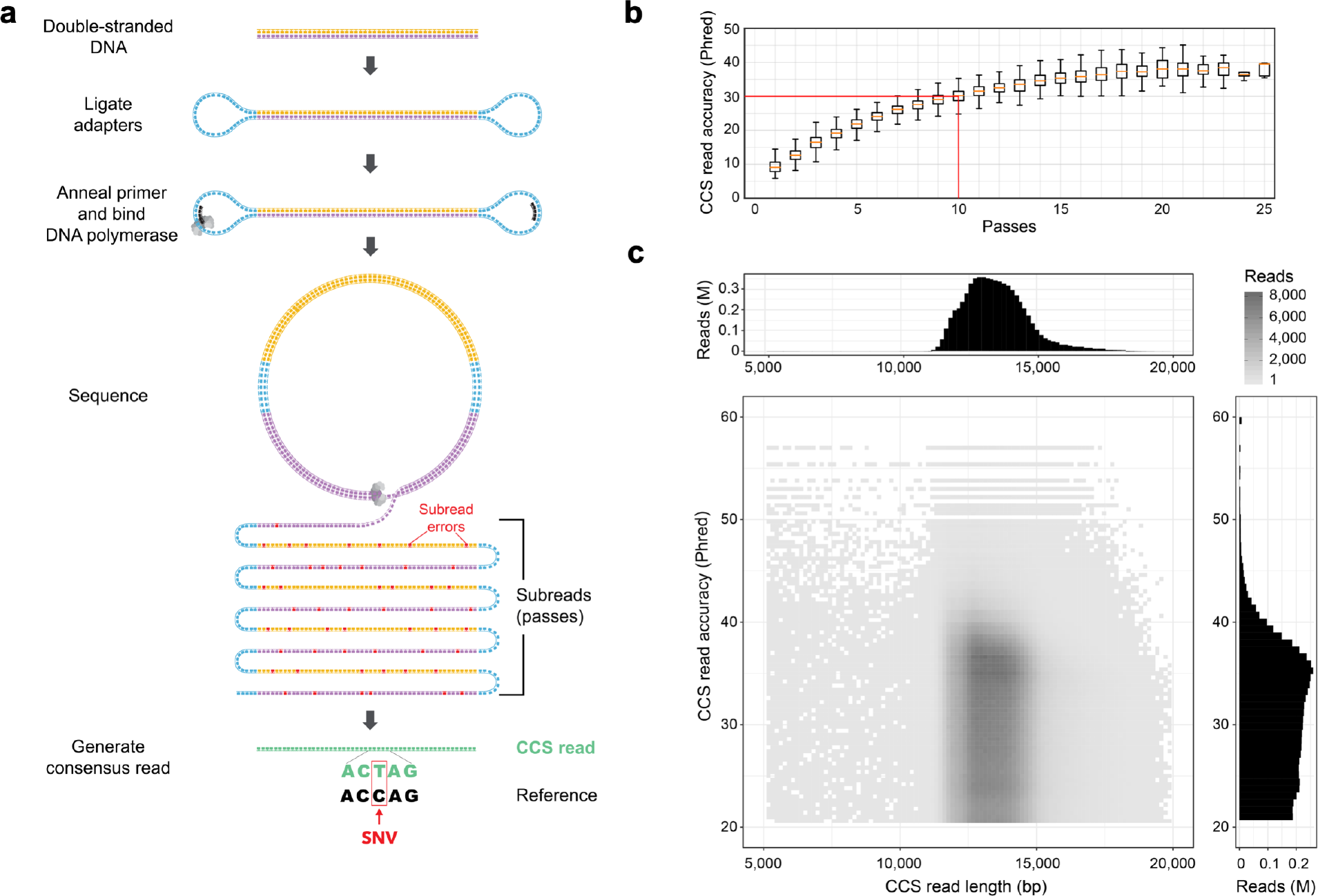
Sequencing HG002 with highly-accurate, long reads. (a) Circular consensus sequencing (CCS) derives a consensus read from multiple passes of a single template molecule, producing accurate reads from noisy individual subreads (passes). (b) Predicted accuracy of CCS reads with different numbers of passes, for sequencing of the human male HG002. At 10 passes, the median read achieves Q30 predicted accuracy. (c) Length and predicted accuracy of CCS reads.

### Quality Evaluation of CCS Reads

To characterize the few residual errors in CCS reads, discordances between the reads and the GIAB HG002 benchmark were tallied. The average read concordance is 99.8%, comparable to the 99.9% concordance of short reads from the Illumina NovaSeq (**Supplementary Table 1**). The large majority of CCS read discordances are indels in homopolymer contexts: 3.4% are mismatches, 4.6% are indels in non-homopolymer contexts, and 92.0% are indels in homopolymers. This equates to a mismatch every 13,048 bp in CCS reads, a non-homopolymer indel every 9,669 bp, and a homopolymer indel every 477 bp (**Supplementary Table 1**). The mismatch rate is 17× lower than reads from the Illumina NovaSeq, while the indel rate is 181× higher (**Supplementary Table 1**).

To confirm independently the high quality of CCS reads, error rates were measured through read-to-read alignments^26^. Consistent with the reference-based methods, the average read accuracy is estimated at 99.8%. A putative large artifact is detected in 0.6% of reads: 0.5% are molecular chimeras, likely due to ligation of DNA fragments during library construction, 0.1% contain a “low quality” run of bases, anecdotally in microsatellites, and 0.03% have a missing SMRTbell adapter on one end. Overall, the read-to-read comparison supports the predicted quality of the CCS reads.

### Increased Mappability of CCS Reads

To evaluate increases in mappability with long reads, the 13.5 kb CCS reads and a coverage-matched number of 2×250 bp NGS short reads were mapped to GRCh37. A genomic position was considered to be mappable if it is covered by least ten reads. At the highest reported mapping quality (60), 97.5% of the non-gap GRCh37 is mappable with 13.5 kb CCS reads, while 94.8% is mappable with NGS short reads (**Figure 2a**).

**Figure 2.**
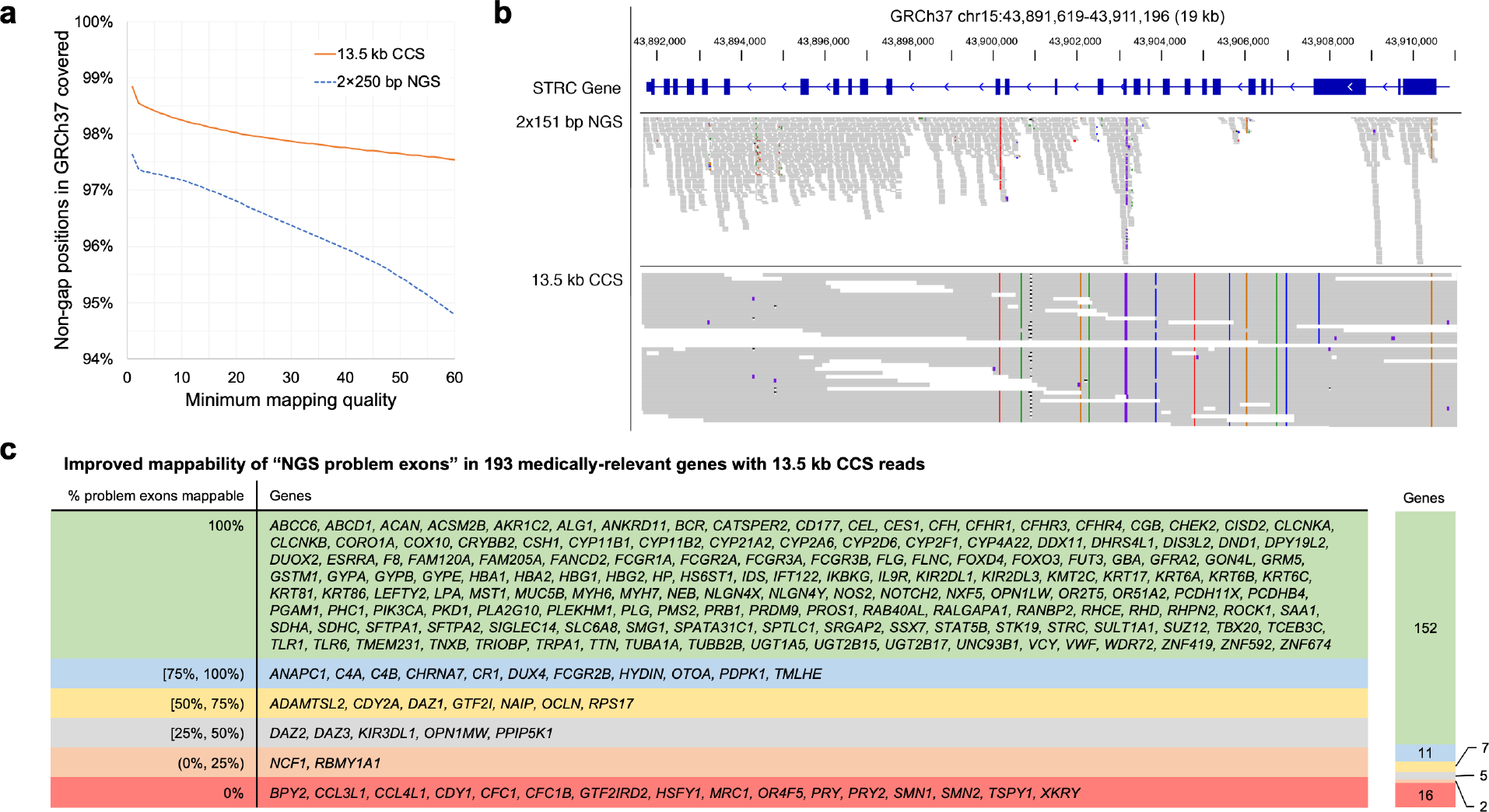
Mappability of the human genome with CCS reads. (a) Percentage of the non-gap GRCh37 human genome covered by at least 10 reads from 28-fold coverage NGS (2×250 bp) and CCS (13.5 kb) datasets at different mapping quality thresholds. (b) Coverage of the congenital deafness gene *STRC* in HG002 with 2×151 bp NGS (NovaSeq) reads and 13.5 kb CCS reads at a mapping quality threshold of 10. (c) Improvement in mappability with 13.5 kb CCS reads for 193 human genes previously reported as medically-relevant and problematic to map with NGS reads^27^.

The additional regions that are now accessible with longer CCS reads include numerous medically-relevant genes which have been previously reported as recalcitrant to NGS sequencing^27^. Of the 193 reported medically-relevant genes with at least one NGS problem exon, 152 are fully mappable with the CCS reads, including *CYP2D6*, *GBA*, *PMS2*, and *STRC* (**Figure 2b-c**).

The 13.5 kb CCS reads also resolve complex regions, like the HLA class 1 and 2 genes, which are fully phased and typed to four-field resolution^28^ (**Supplementary Figure 3**).

### Small Variant Detection with CCS Reads

GATK^29^ was used to call SNVs and small indels with CCS reads. Evaluated against the GIAB benchmark^25^, precision for SNVs is 99.468% and recall is 99.559%. For indels, precision is 78.977% and recall is 81.248%. While GATK performance with CCS reads is comparable to NGS for SNVs, it is lower for indels (**Table 1**). Unlike NGS read errors, which are mostly mismatches, CCS read errors are mostly indels (**Supplementary Table 1**), contributing to the low indel precision and recall of GATK for CCS reads.

**Table 1.** Performance of small variant calling with CCS reads. Precision, recall, and F1 of small variant calling measured against the Genome in a Bottle v3.3.2 benchmark using hap.py. **Bold** indicates the highest value in each column. <u>Underline</u> indicates a value higher than the GATK HaplotypeCaller run on 30-fold Illumina NovaSeq reads. Coverage is 28-fold for PacBio CCS and 30-fold for Illumina NovaSeq. Rows are sorted (“^”) based on F1 for SNVs.

| Platform | Variant caller (training model) | SNVs |  |  | Indels |  |  |
| --- | --- | --- | --- | --- | --- | --- | --- |
|  |  | Precision | Recall | F1 ^ | Precision | Recall | F1 |
| Illumina (NovaSeq) | DeepVariant (Illumina model) | **<u>99.960%</u>** | <u>99.940%</u> | **<u>99.950%</u>** | **<u>99.633%</u>** | **<u>99.413%</u>** | **<u>99.523%</u>** |
| PacBio (CCS) | DeepVariant (CCS model) | <u>99.914%</u> | <u>99.959%</u> | <u>99.936%</u> | 96.901% | 95.980% | 96.438% |
| PacBio (CCS) | DeepVariant (haplotype-sorted CCS model) | <u>99.904%</u> | **<u>99.963%</u>** | <u>99.934%</u> | 97.835% | 97.141% | 97.486% |
| Illumina (NovaSeq) | GATK HaplotypeCaller (no filter) | 99.852% | 99.910% | 99.881% | 99.371% | 99.156% | 99.264% |
| PacBio (CCS) | GATK HaplotypeCaller (hard filter) | 99.468% | 99.559% | 99.513% | 78.977% | 81.248% | 80.097% |

Variant callers based on deep learning have an inherent ability to adapt to the error profiles of new data types. To evaluate variant calling with a deep learning framework, Google DeepVariant^30^ was used to call SNVs and indels from CCS reads. Using a model trained on Illumina reads, precision is 99.533% and recall is 99.793% for SNVs, and precision is 23.991% and recall is 81.692% for indels (**Supplementary Table 2**).

Training a model on CCS reads provides a large boost in precision and recall for both SNVs and indels. For SNVs, DeepVariant achieves precision of 99.914% and recall of 99.959%. For indels, DeepVariant achieves 96.901% precision and 95.980% recall (**Figure 3a**, **Table 1**). Most discordant indels (90.33%) occur in homopolymer runs, matching the most common discordance in CCS reads (**Supplementary Figure 4**). The callset includes 1,969 SNVs and 62 indels in exons of medically-relevant genes previously reported as recalcitrant to NGS sequencing.

**Figure 3.**
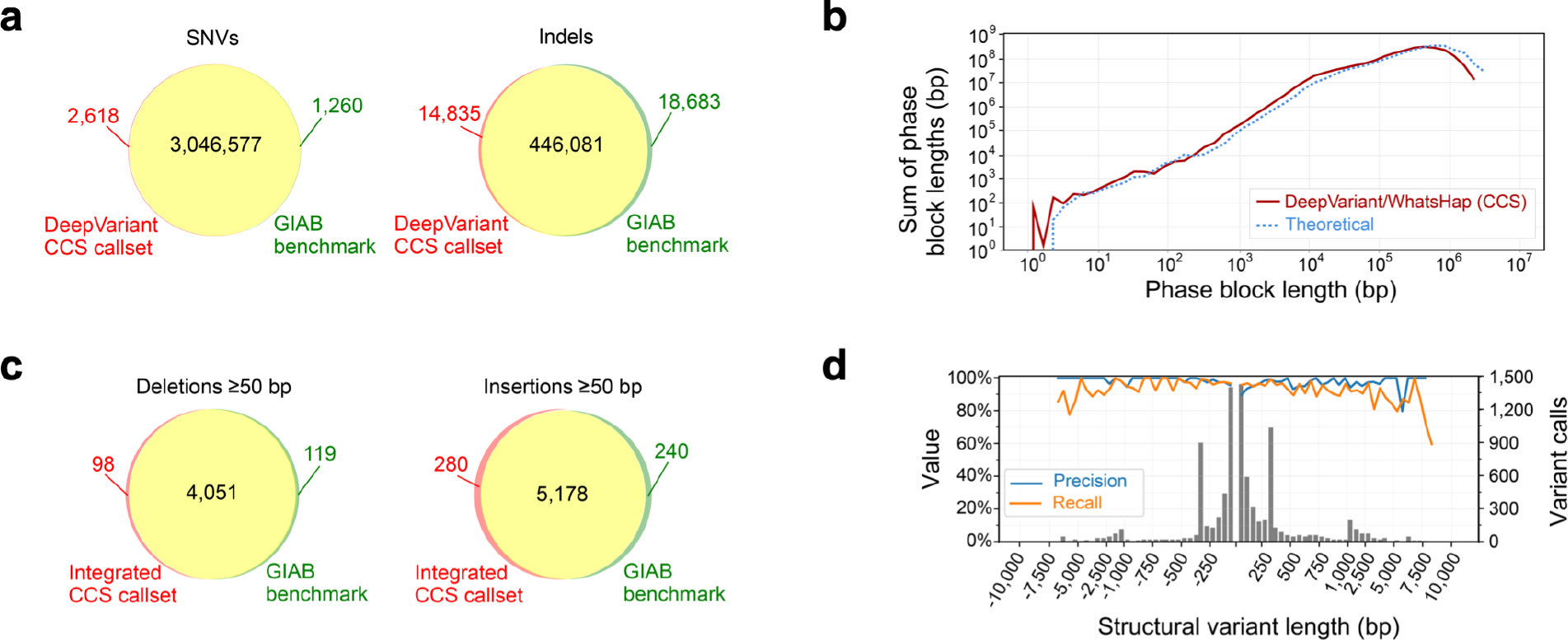
Variant calling and phasing with CCS reads. (a) Agreement of DeepVariant SNV and indel calls with Genome in a Bottle v3.3.2 benchmark measured with hap.py. (b) Phasing of heterozygous DeepVariant variant calls with WhatsHap, compared to theoretical phasing of HG002 with 13.5 kb reads. (c) Agreement of integrated CCS structural variant calls with the Genome in a Bottle v0.6 structural variant benchmark measured with Truvari, (d) by variant length. Negative length indicates a deletion; positive length indicates an insertion. The histogram bin size is 50 bp for variants shorter than 1 kb, and 500 bp for variants >1 kb.

### Phasing Small Variants with CCS Reads

To determine whether CCS reads could provide both highly-accurate variant calls and long-range information needed to generate haplotypes, we used WhatsHap^31^ to phase the DeepVariant variant calls. Nearly all (99.64%) autosomal heterozygous variants were phased into 19,215 blocks with an N50 of 206 kb (**Supplementary Table 3**). The phase block length distribution closely matches the theoretical limit estimated by creating breaks between variants that are separated by more than the average CCS read length of 13.5 kb. This suggests that the phase block length is limited by read length and the amount of variation in HG002, not by coverage or the quality of the variant calls (**Figure 3b**, **Supplementary Figure 5**). Evaluated against the GIAB benchmark phase set, the switch error rate is 0.37% and the Hamming error rate is 1.91% (**Supplementary Table 3**).

### Improving Small Variant Detection with Haplotype Phasing

GATK and DeepVariant do not directly incorporate long-range haplotype phase information when calling variants. To evaluate whether phase information improves results, CCS reads were haplotype-tagged based on trio-phased variants from GIAB (which tags 84.55% of reads) and a DeepVariant model was then trained on reads passed in haplotype-sorted order. The haplotype-sorted model performs similarly to the original DeepVariant CCS model for SNVs but provides a large improvement for indels achieving precision of 97.835% and recall of 97.141% (**Table 1**).

### Structural Variant Detection with CCS Reads

Insertion and deletion structural variants ≥50 bp were called using two read mapping-based tools, pbsv (https://github.com/PacificBiosciences/pbsv) and Sniffles^32^. The callsets show similar precision (>94%) and recall (>91%) against the GIAB benchmark (**Supplementary Table 4**). Precision is consistent across variant length, but recall is lower for variants ≥3kb (**Supplementary Figure 6a-b**). To increase recall for larger variants, haplotype-resolved *de novo* assemblies were analyzed with paftools^33^ (see “*De Novo* Assembly of CCS Reads”), with precision >93% and recall >89% (**Supplementary Figure 6c-d, Supplementary Table 4**).

An integrated callset includes 8,432 deletions and 12,091 insertions. Precision is 96.13% and recall is 95.99% (**Figure 3c**, **Supplementary Table 4**), with similar performance for insertions as deletions and for variants <1kb as ≥1kb (**Figure 3d**), indicating the complementarity of mapping- and assembly-based structural variant calling. The callset has 163 deletions and 143 insertions that intersect exons.

For comparison, structural variants were called in Illumina 2×250 bp short reads (with Manta^34^ and Delly^35^) and 10X Genomics linked reads (with LongRanger^36^) available from GIAB^24^. The Manta callset has precision of 85.34% and recall of 55.88%, with much worse recall for insertions (39.65%) than deletions (76.90%). The LongRanger callset has precision of 83.79% and recall of 39.83%, again with worse recall for insertions (16.41%) than deletions (70.18%). A callset from paftools run on a linked-read SuperNova assembly has precision of 64.52% and recall of 52.74% (**Supplementary Table 4**, **Supplementary Figure 6e-h**). All considered short- and linked-read callsets have worse performance than all CCS callsets in both precision and recall.

### *De Novo* Assembly of CCS Reads

Three different algorithms – FALCON^37^, Canu^38^, and wtdbg2 (https://github.com/ruanjue/wtdbg2) – were used to assemble the full CCS read set, which is a mix of paternal and maternal reads. By skipping the initial read-to-read error correction step, the algorithms completed 10-100× faster than is typical for long-read assemblies^21^ (**Supplementary Table 5**). All assemblies have high contiguity with a contig N50 from 15.43 to 28.95 Mb. The total assembly size is near the expected human genome size for FALCON and wtdbg2. The Canu assembly has a total genome size of 3.42 Gb, larger than the expected haploid human genome, because it resolves some heterozygous alleles into separate contigs (**Table 2**, **Supplementary Figure 7**).

**Table 2.** Statistics for *de novo* assembly of CCS reads. The “mixed” haplotype assemblies use all reads. The “maternal” and “paternal” assemblies use parent-specific reads from trio binning plus unassigned reads. HG002 concordance is measured against the Genome in a Bottle benchmark. BUSCO gene completeness uses the Mammalia ODB9 gene set. RefSeq genes is the percentage of genes from Ensembl R94 that are full-length, single-copy in the assembly relative to the full-length, single-copy count for GRCh38. Contigs shorter than 13 kb were excluded from genome size and contiguity measurements; contigs shorter than 100 kb were excluded from the concordance measurement. “*” indicates polishing with Arrow.

| Haplotype | Assembler | Total size (Gb) | Contigs | N50 (Mb) | NG50 (Mb) | Max (Mb) | E-size<sup>53</sup> (Mb) | HG002 concordance (Phred) | BUSCO genes | RefSeq genes |
| --- | --- | --- | --- | --- | --- | --- | --- | --- | --- | --- |
| Mixed | Canu | 3.42 | 18,006 | 22.78 | 25.02 | 108.46 | 30.16 | 31.1 | 92.3% | 93.2% |
| Mixed | FALCON | 2.91 | 2,541 | 28.95 | 24.51 | 110.21 | 38.04 | 25.8 | 87.6% | 97.6% |
| Mixed | wtdbg2 | 2.79 | 1,554 | 15.43 | 12.62 | 84.67 | 22.61 | 44.6 | 94.2% | 96.1% |
| Maternal | Canu* | 3.04 | 5,854 | 18.02 | 17.04 | 48.81 | 19.78 | 47.2 | 94.1% | 98.1% |
| Maternal | FALCON* | 2.80 | 924 | 19.99 | 15.54 | 74.33 | 24.07 | 43.5 | 95.1% | 97.8% |
| Maternal | wtdbg2 | 2.75 | 2,637 | 12.10 | 9.29 | 66.34 | 16.55 | 43.5 | 93.8% | 95.6% |
| Paternal | Canu* | 2.96 | 6,868 | 16.14 | 14.90 | 64.83 | 20.19 | 47.7 | 93.4% | 98.2% |
| Paternal | FALCON* | 2.70 | 1,489 | 16.40 | 14.06 | 95.34 | 25.61 | 43.5 | 93.6% | 97.7% |
| Paternal | wtdbg2 | 2.67 | 1,444 | 13.96 | 10.86 | 50.51 | 15.36 | 42.1 | 92.6% | 95.3% |

Short reads from the parents of HG002 were used to identify k-mers unique to one parent and then partition (“trio bin”) the CCS reads by haplotype^39^. Three different k-mer sizes were evaluated: 21 bp (previously reported for trio binning) and longer k-mers of 51 bp and 91 bp enabled by the accuracy of CCS reads. The 21-mer binning assigns 35.3% of reads to the mother and 33.6% to the father (68.9% binned). The 51-mer binning is more complete at 78.5% binned; using longer 91-mers provides only a small additional gain to 79.2% binned. The 51-mer binning was selected for assembly (**Supplementary Table 6**).

FALCON, Canu, and wtdbg2 were run separately on the paternal and maternal reads, with the unassigned reads included in both sets. All algorithms produce highly contiguous and nearly complete assemblies for the parental genomes, with N50 from 12.10 to 19.99 Mb and genome size from 2.67 to 3.04 Gb (**Table 2**). From 95.3% to 98.2% of human genes are identified as single-copy in each parental assembly (**Table 2**). Assembly-based structural variant calls have high precision and recall, suggesting few large-scale mis-assemblies (**Supplementary Table 4**). Furthermore, analysis of the phase-consistency^40^ of maternal and paternal haplotigs shows the assemblies are phased properly (**Supplementary Figure 8**).

All mixed and parental assemblies are high quality with concordance to the HG002 benchmark ranging from Q44-Q48 for polished^41^ and Q26-Q45 for unpolished assemblies (**Table 2**, **Supplementary Table 7**). This greatly exceeds that of previously published and accessioned assemblies at Q40 (6× worse) for PacBio noisy long reads and Q29 (77× worse) for Oxford Nanopore reads with Illumina polishing (**Figure 4a**, **Supplementary Table 7**).

**Figure 4.**
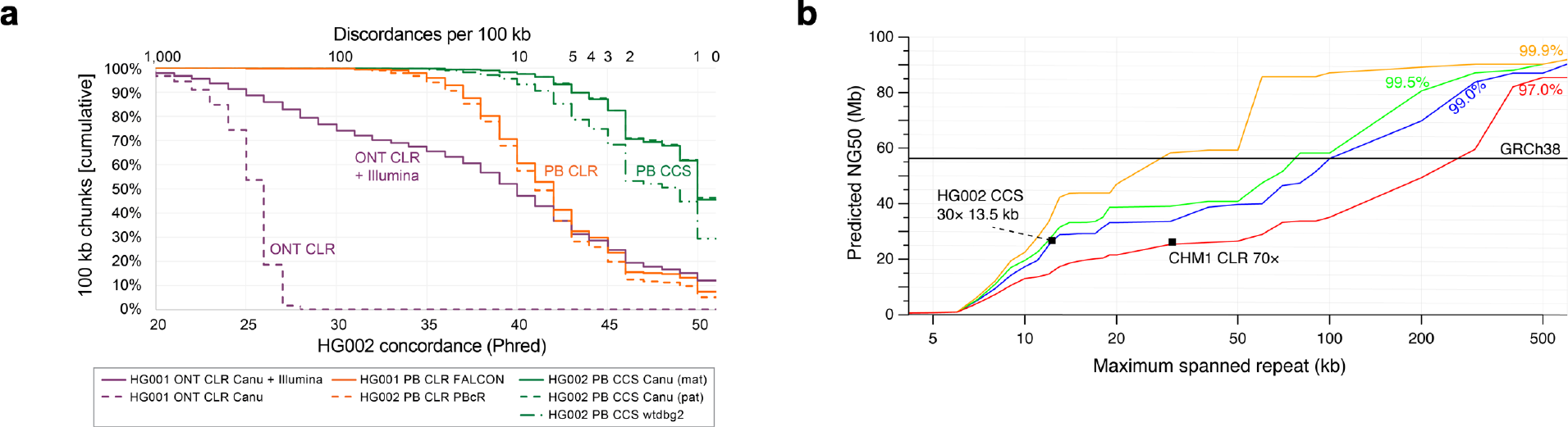
Impact of read accuracy on *de novo* assembly. (a) The concordance of seven assemblies to the Genome in a Bottle HG002 benchmark (**Supplementary Table 7**). Contigs longer than 100 kb were segmented into 100 kb chunks and aligned to GRCh37. Concordance was measured per chunk, and chunks with no discordances were assigned concordance of Q51. PB=PacBio, ONT=Oxford Nanopore, CLR=continuous (“noisy”) long reads. (b) Predicted contiguity of a human assembly based on ability to resolve repeats of different lengths (x-axis) and percent identities (colored lines)^21^. The solid line indicates the contiguity of GRCh38. The 97.0% identity line is representative of CLR assemblies using standard read-to-read error correction. The points show example CCS and CLR^43^ assemblies using Canu. Repeat identity and length are proxies for read accuracy and length.

Large segmental duplications often result in contig breaks in *de novo* assemblies, and assemblies of noisy long reads typically span less than 50 of 175 Mb of segmental duplications in the human genome^15,18,42^. The most contiguous assemblies of CCS reads span over 60 Mb of segmental duplications, a 20% improvement (**Supplementary Table 8**). A model of assembly contiguity based on large repeat resolution suggests that the current assemblies of CCS reads resolve 15 kb repeats of 99 to 99.5% identity (**Figure 4b**).

### Coverage Requirements for Variant Calling and *De Novo* Assembly

To evaluate the depth of CCS read coverage required for variant calling and assembly, we randomly subsampled from the full dataset. For SNVs, precision and recall with DeepVariant remain above 99.5% for coverage down to 15-fold; performance decays steeply below 10-fold (**Supplementary Figure 9a**). For indels, DeepVariant remains comparable to typical NGS performance (>90%) down to 17-fold coverage (**Supplementary Figure 9b**). For structural variants, precision with pbsv is above 95% for all evaluated coverage levels. Recall is above 90% down to 15-fold coverage, and decays steeply below 10-fold (**Supplementary Figure 9c**). For phasing with WhatsHap, the phase block N50 remains above 150 kb down to 10-fold coverage (**Supplementary Figure 9d**). Mixed-haplotype wtdbg2 assemblies have consistent size above 2.7 Gb, contig N50 around 15 Mb, and concordance above Q42 until coverage falls below 15-fold (**Supplementary Figure 9e-g**).

### Revising and Expanding Genome in a Bottle Benchmarks

High-quality callsets from CCS reads provide an opportunity to identify mistakes in the GIAB benchmarks, particularly for structural variants where the benchmark is still in draft form. Sixty small variant and 40 structural variant discrepancies between the GIAB benchmark (small variant v3.3.2, structural variant v0.6) and the CCS callsets (DeepVariant haplotype-sorted, structural variant integrated) were selected for manual curation. Selected variants were spread across variant types, discrepancy types, and both inside and outside homopolymers and tandem repeats.

For small variants, 29 of 31 discrepancies in homopolymers were classified as correct in the benchmark. Outside of homopolymers, 19 of 29 were classified as errors in the benchmark. Most of these benchmark errors (13 of 19) are true variants in L1 elements called homozygous reference in GIAB (**Supplementary Figure 10a-b**, **Supplementary Table 9**). The identified benchmark errors overlap with putative errors in a DeepVariant Illumina whole genome case study (https://github.com/google/deepvariant). Of 745 putative false positive SNVs in the case study, 344 agree with the CCS callset, with 282 (82.0%) falling within large interspersed repeats. Fewer of the false negative SNVs (8%), false negative indels (25%), and false positive indels (19%) from the case study agree with the CCS callset. Extrapolating from manual curation, we estimate that 2,434 (1,313-2,611; 95% confidence interval) errors in the current GIAB benchmark could be corrected using the CCS reads.

For structural variants, curator classification was unclear for 11 of 40 discrepancies, typically because of tandem repeat structure that permits multiple representations of a variant. For the remainder, 15 of 16 false negative discrepancies were classified as correct in the benchmark. However, for false positive discrepancies, 11 of 13 were classified as errors in the benchmark (**Supplementary Figure 10c-d**, **Supplementary Table 10**). This suggests that the GIAB structural variant benchmark set is precise but incomplete.

The high-quality CCS callsets also provide an opportunity to expand the benchmarks into repetitive and highly polymorphic regions that have been difficult to characterize with confidence using short reads. Adding the CCS DeepVariant callset to the existing GIAB small variant integration pipeline would expand the benchmark regions by up to 1.3% and 418,875 variants (210,184 SNVs and 208,691 indels). For structural variants, only 9,232 of 18,832 autosomal variant calls overlap benchmark regions, which means that the number of variants in the benchmark would more than double if all CCS variants calls were incorporated.

## Discussion

We present a protocol for producing highly-accurate long reads using circular consensus sequencing (CCS) on the PacBio Sequel System. We apply the protocol to sequence the human HG002 to 28-fold coverage with average read length of 13.5 kb and an average read accuracy of 99.8%. We analyze the CCS reads to call SNVs, indels, and structural variants; to phase variants into haplotype blocks; and to *de novo* assemble the HG002 genome.

The CCS performance for SNV and indel calling rivals that of the commonly-used pairing of BWA and GATK on 30-fold short-read coverage. Interestingly, though the overall accuracy of CCS reads is similar to short reads, direct application of the GATK pipeline to CCS reads produces inferior results, especially for indels. The major residual error in CCS reads – indels in homopolymers – is not as frequent in short reads. We suspect that the current GATK, which was designed for short reads, does not properly model the CCS error profile, and thus performance lags for indels. This is supported by results with DeepVariant. When a DeepVariant model trained on Illumina reads is run on CCS reads, the performance is poor for indels. When DeepVariant is trained on CCS reads, performance improves dramatically. As more CCS datasets are made available, both model-based callers like GATK and learning-based callers like DeepVariant will have the opportunity to improve on the performance reported here, including by incorporating haplotype phase information and evaluating and training against updated GIAB benchmarks that correct errors with CCS reads. Further, advances in sequencing chemistry or consensus base calling (such as the application of deep learning) that reduce the residual indel errors in CCS reads also could improve variant calling performance.

Structural variant calling and *de novo* genome assembly with CCS reads match or exceed that reported for noisy long reads. The CCS reads have an advantage of high accuracy, which eliminates the need for read correction, allows more stringent criteria to be used in variant calling or read overlapping, and ultimately produces more accurate assemblies and variant calls. Noisy long reads have an advantage of longer maximum read length, but increased accuracy of CCS reads compensates for the length required for highly contiguous assembly. Modeling (**Figure 4b**) suggests modest advances in accuracy (to 99.9%) at 15 kb read length would double the current contiguity, which already matches the best published *de novo* assemblies^43^.

The CCS read approach alleviates some other challenges of long-read sequencing. First, aiming for fragments in the 10-20 kb size range relaxes the need to isolate ultra-long genomic DNA. Second, increased accuracy allows for more stringent alignment and overlap comparisons, greatly reducing the compute time and cost while improving assembly results by recognizing fine-grained repeat and haplotype phase information. Third, familiar tools like GATK that were developed for accurate short reads are readily applied to CCS reads.

Variant calling and assembly with CCS reads perform well down to 15-fold coverage, which offsets the current reduced throughput per run compared to noisy long reads. Ultimately more unique reads per run (5-10× is expected) and increased read length to allow larger fragment sizes will facilitate rapid, population-scale analysis of full genomes with CCS reads to improve human health.

## Methods

### CCS Library Preparation

PacBio library preparation and sequencing was performed on the human reference genome sample HG002 obtained from NIST. Genomic DNA was sheared using the Megaruptor^®^ from Diagenode with a long hydropore cartridge and a 20 kb shearing protocol. Prior to library preparation, the size distribution of the sheared DNA was characterized on the Agilent 2100 BioAnalyzer System using the DNA 12000 kit. A sequencing library was constructed from this sheared genomic DNA using the SMRTbell™ Template Prep Kit v 1.0 (Pacific Biosciences Ref. No. 100-259-100). In order to tighten the size distribution of the SMRTbell™ library, the sample was separated into 3 kb fractions using the SageELF System from Sage Science. Fractions having the desired size distribution ranges were identified on the Agilent 2100 BioAnalyzer using the DNA 12000 kit (**Supplementary Figure 1**). The fraction centered at 15 kb was used for sequencing.

### Sequencing

Sequencing reactions were performed on the PacBio Sequel System with the Sequel Sequencing Kit 3.0 chemistry. The samples were pre-extended without exposure to illumination for 12 hours to enable the polymerase enzymes to transition into the highly processive strand-displacing state and sequencing data was collected for 24 hours to ensure suitably long read lengths. Average subread yield was 35.3±6.2 Gb over 39 SMRT Cells with average polymerase read length of 100.0 kb.

Consensus reads (“CCS reads”) were generated using the ccs software version 3.0.0 (https://github.com/pacificbiosciences/unanimity/) with `--minPasses 3 -- minPredictedAccuracy 0.99 --maxLength 21000`. Average run time is 3,035 CPU core hours per SMRT Cell (118,365 total). The total CCS read yield was 89 Gb (2.3±0.4 Gb per SMRT Cell), with read length of 13.5±1.2 kb.

### Read Mapping

Reads were mapped to the GRCh37 human reference genome, specifically the hs37d5 build from the 1000 Genome Project^44^. CCS reads were mapped with pbmm2 version 0.10.0 (https://github.com/PacificBiosciences/pbmm2) with `--preset CCS`. NGS reads were mapped with minimap2^33^ version 2.14-r883 with `-x sr`.

### Measuring HG002 Concordance

To measure concordance to HG002, alignments to GRCh37 were evaluated at positions within GIAB v3.3.2 benchmark high-confidence regions that have no high-confidence variant call^25^. `Concordance = M/(M+X+D+I)` where M is the number of matches, X is the number of mismatches, D is the number of deletion basepairs, and I is the number of insertion basepairs. Phred = −10*log_10_(1−Concordance). Reads with perfect concordance are assigned a Phred score of 1+log_10_(ReadLength).

A deleted basepair is considered a homopolymer deletion when it matches the preceding or following basepair in the reference genome. An insertion is considered a homopolymer insertion when the basepairs of the insertion are identical and match either the preceding or following basepair in the reference genome.

### Coverage by [GC] content

To measure coverage by local [GC] content, bedtools^45^ version 2.27.1 was used to divide the GRCh37 reference genome into 500 bp windows (`bedtools makewindows-w 500`) and then to calculate the [GC] content (`bedtools nuc`) and average coverage (`bedtools coverage -mean`) of each window.

### Reference-independent Quality Evaluation

The Dazzler suite (https://dazzlerblog.wordpress.com/) was used to evaluate the accuracy of the CCS reads without relying on a reference genome. Briefly, daligner^26^ was used to produce all local alignments longer than 1 kb between pairs of CCS reads. Each CCS subject read was partitioned into 100 bp panels, within which its coverage by and concordance to aligned target reads was calculated. Panels with a concordance in the worst 0.1% were considered low quality. Abrupt breaks in read-to-read alignments were used to estimate library artifacts like chimeric molecules and missing adapters.

### Mappability of CCS and NGS Reads

To compare with the mappability of 13.5 kb CCS reads, a coverage-matched (89 Gb) set of 2×250bp Illumina HiSeq 2500 reads for HG002 were obtained from GIAB^24^ and mapped to GRCh37 with minimap2^33^ version 2.14-r883 with `-x sr`.

A genome position is considered mappable if it is covered by alignments for at least ten reads at a specified mapping quality or higher, which was evaluated using `bedtools bamtobed` and `bedtools genomecov -bga`. Gaps (“N” basepairs in the reference) were excluded.

Previously-reported NGS problem exons^27^ were considered mappable if every basepair in the exon is covered by a read at mapping quality of 60.

### HLA Typing

The *HLA-A* and *HLA-DPA1* genes were typed by comparing the sequence of CCS reads that span the genes to entries in the IMGT database^46^ version 3.19.0.

### Small Variant Detection and Benchmarking

To develop a workflow for calling variants in CCS reads with GATK^29^ HaplotypeCaller v4.0.6.0, different values of the HaplotypeCaller parameter `--pcr-indel-model` and VariantFiltration parameter `--filter-expression` were considered to maximize SNV and indel F1 without excessive complication, starting from the GATK best practices for hard filtering. In the end, HaplotypeCaller was run on reads with a minimum mapping quality of 60 using allele-specific annotations (`--annotation-group AS_StandardAnnotation`) and `--pcr_indel_model AGGRESSIVE`. Autosomes and the pseudo-autosomal regions (PARs) on chromosome X were called with `--ploidy 2`; chromosome Y and the non-PAR regions of chromosome X were called with `--ploidy 1`. Multi-allelic variant sites were split into separate entries for filtration with a custom script (https://gist.github.com/williamrowell/16cd89fcb23ab9f11a7bd387c308d29d). SNVs were filtered using GATK VariantFiltration with `--filter_expression of AS_QD < 2.0` for SNVs and indels longer than 1bp, and `AS_QD < 5.0` for 1 bp indels. A similar pipeline was used to call variants in coverage-matched 2×151 bp Illumina NovaSeq reads with a few differences: a minimum mapping quality of 20, `--pcr-indel-model NONE, --standard_min_confidence_threshold_for_calling 2.0`, and no variant filtration.

A Google DeepVariant model for CCS reads was generated as previously reported^30^ using DeepVariant version 0.7.1. Briefly, models were trained using CCS reads for chromosomes 1-19 and the HG002 GIAB v3.3.2 benchmark. A single model was selected based on performance in chromosomes 21 and 22 to avoid overfitting. Neither training nor model selection considers chromosome 20, which is available for accuracy evaluations. To support long reads, local reassembly is disabled for DeepVariant with CCS reads. The wgs_standard model version 0.7.1 was used to call variants in NovaSeq reads and to apply a model trained on Illumina reads to CCS reads.

To incorporate long-range haplotype information, DeepVariant was modified to produce pileups with reads sorted by the BAM haplotype (“HP”) tag. Haplotype information was added to the pbmm2 CCS alignments using WhatsHap v0.17 (`whatshap haplotag`) with the trio-phased variant calls from GIAB (ftp://ftp-trace.ncbi.nlm.nih.gov/giab/ftp/data/AshkenazimTrio/analysis/NIST_MPI_whatshap_08232018/RTG.hg19.10x.trio-whatshap.vcf.gz or https://bit.ly/2R73grR). A new DeepVariant model then was trained as described above.

Small variant callsets were benchmarked against the GIAB v3.3.2 HG002 set^25^ by vcfeval^47^ (https://github.com/RealTimeGenomics/rtg-tools) with no partial credit run through hap.py version 0.3.10 (https://github.com/Illumina/hap.py). Only PASS calls were considered.

### Phasing Small Variants

Small variant calls were phased using WhatsHap v0.17 (`whatshap phase`). The number of switch and Hamming errors was computed against trio-phased variant calls from GIAB using `whatshap compare`.

To model the phase blocks achievable with a given read length, cuts were introduced between heterozygous variants in the GIAB trio-phased variant callset that are separated by more than the read length, which effectively assumes that adjacent heterozygous variants separated by less than the read length can be phased.

### Structural Variant Detection

pbsv version 2.1.0 (https://github.com/PacificBiosciences/pbsv) was run on pbmm2 CCS read alignments. The `pbsv discover` stage was run separately per chromosome with tandem repeat annotations (https://github.com/PacificBiosciences/pbsv/tree/master/annotations) passed with `--tandem-repeats`. The `pbsv call` stage was run on the full genome.

Sniffles version 1.0.10 was run on pbmm2 CCS reads alignments with `-s 3 --skip_parameter_estimation` and with the variant sequence obtained from reads.

Structural variants in the maternal and paternal Canu and FALCON assemblies from CCS reads (see “*De novo* Assembly”) were called using a previously described workflow^48^. Briefly, contigs were mapped to GRCh37 using `minimap2 --paf-no-hit --cxasm5 --cs -r2k`; variants were called with paftools.js call^33^; maternal and paternal variants were concatenated; and indel calls at least 30 bp were retained.

An integrated callset was produced from the pbsv, Sniffles, and paftools/Canu callsets using SURVIVOR^49^ and custom scripts. Two calls were considered supporting if the calls had the same structural variation type, a start position with 1 kb, and a difference in length less than 5%. One call from each matching set was retained with precedence given to pbsv, then Sniffles, and then paftools. Because pbsv and Sniffles have poor sensitivity for calls larger than 1 kb, all non-matched calls from paftools that are larger than 1 kb were retained.

NovoAlign (http://www.novocraft.com) alignments to GRCh37 of 300-fold coverage of HG002 with 2×250bp Illumina HiSeq 2500 reads were obtained from GIAB. Structural variants were called with Manta^34^ version 1.4.0 with all coverage, and Delly^35^ version 0.7.6 with coverage subsampled to 30-fold using `samtools view -b -s 0.1`.

Structural variant callsets on 10X Genomics reads from LongRanger version 2.2 were obtained from GIAB (ftp://ftp-trace.ncbi.nlm.nih.gov/giab/ftp/data/AshkenazimTrio/analysis/10XGenomics_ChromiumGenome_LongRanger2.2_Supernova2.0.1_04122018/ or https://bit.ly/2Mtj084). Insertion and deletion variants at least 30 bp were combined from the sequence-resolved indels and large deletion calls (NA24385_LongRanger_snpindel.vcf.gz, NA24385_LongRanger_sv_deletions.vcf.gz). Another callset was produced using paftools on the diploid Supernova 2.0.1 assembly as described above.

Structural variant callsets were benchmarked against the GIAB v0.6 HG002 structural variant set (ftp://ftp-trace.ncbi.nlm.nih.gov/giab/ftp/data/AshkenazimTrio/analysis/NIST_SVs_Integration_v0.6 or https://bit.ly/2T7iLBX) using Truvari (https://github.com/spiralgenetics/truvari) commit 600b4ed7 modified to allow a single variant in the benchmark set to support multiple variants in the callset. Truvari was run with `-r 1000 -p 0.01 -- multimatch --includebed HG002_SVs_Tier1_v0.6.bed -c HG002_SVs_Tier1_v0.6.vcf.gz`. The `-p 0` option was used to disable sequence checks for callsets that report symbolic alleles instead of sequence-resolved calls (LongRanger, Delly).

### *De novo* Assembly

Mixed haplotype assemblies were produced using all CCS reads. Canu^38^ version 1.7.1 was run with `-p asm genomeSize=3.1g correctedErrorRate=0.015 ovlMerThreshold=75 batOptions="-eg 0.01 -eM 0.01 -dg 6 -db 6 -dr 1 -ca 50 -cp 5" -pacbio-corrected`. FALCON^37^ kit version 1.2.0 was run with `ovlp_HPCdaligner_option = -v -B128 -M24 -k24 -h1024 -e.97 -l2500 -s100, ovlp_DBsplit_option = -s400`, and `overlap_filtering_setting = --max-diff 90 --max-cov 120 --min-cov 2`. Wtdbg2 (https://github.com/ruanjue/wtdbg2) version 2.2 was run with `-k 0 -p 21 -AS 4 -s 0.5 -e 2 -K 0.05` and followed by wtdbg2-cns.

CCS reads from HG002 were “trio binned” as maternal, paternal, or unassigned as previously described^39^. Briefly, 2×250 bp Illumina HiSeq 2500 reads for the father (HG003/NA24149) and mother (HG004/NA24143) of HG002 were obtained from GIAB. Sequence k-mers unique to the mother or father were identified and used to categorize CCS reads (https://github.com/skoren/triobinningScripts), using k-mer size of 21, 51, and 91 and excluding k-mers that occur 25 times or fewer. The maternal and unassigned reads were used for the “maternal” assemblies; paternal and unassigned reads were used for the “paternal” assemblies.

The maternal and paternal assemblies were generated with Canu and wtdbg2 using the same software version and same options as for the mixed haplotype assembly. For the maternal and paternal assemblies, FALCON version 0.7 was run with length_cutoff_pr = 2000, ovlp_HPCdaligner_option = -k24 -e.95 - s100 -l1000 -h600 -mdust -mrep8 -mtan -M21, ovlp_DBsplit_option = -x2000 -s400, falcon_sense_option = --min_idt 0.70 --min_cov 4 - -max_n_read 200, and `overlap_filtering_setting = --max_diff 40 -- max_cov 80 --min_cov 2 --min_len 500`.

The maternal and paternal Canu assemblies were polished with Arrow version 2.2.2 run through ArrowGrid (https://github.com/skoren/ArrowGrid) using subreads that correspond to the CCS reads used for each assembly. The maternal and paternal FALCON assemblies were polished with Arrow version 2.2.2 using all subreads.

### Assembly Evaluation

For each assembly, contigs were broken into 100 kb chunks with remainders shorter than 100 kb ignored. The chunks were aligned to GRCh37 using `minimap2 --eqx - x asm5`, and primary alignments that span at least 50 kb in the reference at higher than 50% identity were retained. The concordance of each chunk was evaluated just as for CCS reads (see “Measuring HG002 Concordance”). The overall assembly concordance was calculated as the average concordance of the 100 kb chunks.

Gene completeness was measured using BUSCO^50^ version 3.0.2 using the Mammalia ODB9 gene set. The single plus duplicated gene count in the BUSCO summary is reported. For a human-specific measure of completeness, we calculated the fraction of single-copy human genes that remain single-copy in each assembly. The human transcript sequences from ENSEMBL^51^ build r94 were mapped to each assembly with `minimap2 -cx splice -B 4 -O 4,34 -C9 -uf --cs` and evaluated with `paftools.js asmgene -i 0.98`, which retains the longest of overlapping transcripts, and counts a transcript hit if 99% of the transcript sequence maps at 98% identity or higher. A single-copy transcript has exactly one hit. Counts are normalized to the number of transcripts that are considered single-copy by these criteria in GRCh38 (GCA_000001405.15).

To measure the number of segmental duplications spanned by each assembly, the assemblies were processed with segDupPlots^42^ (https://github.com/mvollger/segDupPlots), which maps contigs to GRCh38 and considers a segmental duplication to be spanned by the assembly if a contig alignment extends through the segmental duplication with at least 50 kb on each flank.

### Model of Assembly Contiguity

To predict assembly contiguity at different read lengths and read accuracies, a previously described model^21^ was updated with improvements for high-accuracy reads. Briefly, all repeat annotations for GRCh38 were downloaded from the UCSC Genome Browser. Repeat identity was defined as by each track except for: the nested repeat track where identity was 50+50*score/1000, RepeatMasker where identity was 1-((mismatches + deleted + inserted)/1000), and microsat and windowmasker/sdust which does not define identity and thus was treated as 100%. Gaps were included as 100% identity repeats. Additional repeats were added from self-matches using MashMap^52^ (https://github.com/marbl/MashMap).

The assembly contiguity was predicted based on the ability to resolve repeats. At a given percent identity, repeats below that identity were excluded and remaining repeats separate by 15 bp or fewer were merged. Then, cuts were introduced at each at repeats of each given length, and assembly NG50 was calculated assuming that contigs end at each cut.

### Coverage Titration

To evaluate the performance of variant calling and assembly at different coverage levels, CCS reads were downsampled from the 28-fold dataset and processed. For small variant calling, alignments were subsampled in DeepVariant version 0.7.1 from 4% to 100% in steps of 3%. Variants were called on each subsample using the DeepVariant CCS model. Precision and recall for SNVs and indels were evaluated with hap.py as described above (see “Small Variant Detection and Benchmarking”). For phasing, alignments were subsampled (`samtools view -s`) at rates from 10% to 100% in steps of 10%. The DeepVariant callset from the full 28-fold coverage data was phased using WhatsHap v0.17 (`whatshap phase`) with the subsampled alignments. For structural variants, alignments were subsampled (`samtools view -s`) at rates from 10% to 100% in steps of 10%. Variants were called on the subsampled alignments with pbsv version 2.1.0 and benchmarked with Truvari as described above (see “Structural Variant Detection”). For assembly, reads were subsampled at rates from 10% to 100% in steps of 10%. Sampling was performed based on read name (10% sample is reads that end in 0, 20% is reads that end in 0-1, and so on). Assembly of subsamples reads was performed with wtdbg2 version 2.2 and benchmarked as described above (see “*De novo* Assembly” and “Assembly Evaluation”).

### Revising and Expanding Genome in a Bottle Benchmarks

Discrepancies between the GIAB v3.3.2 small variant benchmark and the DeepVariant callset from haplotype-sorted CCS reads were identified with vcfeval and hap.py. Discrepancies between the GIAB v0.6 structural variant benchmark and the integrated structural variant callset from CCS reads were identified with Truvari. A sample of 60 small variant and 40 structural variant discrepancies were selected for manual curation by random sampling across discrepancy types (false positive, false negative, genotype difference), variant types (SNV, indel, insertion structural variant, and deletion structural variant), both inside and outside homopolymers and tandem repeats. Curators evaluated variants in IGV along with alignments of CCS reads, 10X Genomics reads, Illumina short reads, and Illumina reads from a 6 kb mate pair library, all obtained from GIAB. The benchmark error rate was estimated by variant type and discrepancy type and used to extrapolate from the sample to the number of errors in the full GIAB benchmark. Confidence intervals were calculated assuming a binomial distribution.

## Data Availability

CCS reads and alignments to GRCh37 are available at ftp://ftp-trace.ncbi.nlm.nih.gov/giab/ftp/data/AshkenazimTrio/HG002_NA24385_son/PacBio_CCS_15kb/ or https://bit.ly/2RW1b3I.

## Author Contributions

AMW, DRR, MWH, and PP designed the study. DRR and PP developed the sample preparation protocol and performed sample preparation. DRR, PP, and YQ performed sequencing. AC, AK, CSC, MAD, and PC adapted the algorithms and implementation of DeepVariant. AC, AF, AK, AMP, AMW, AT, CSC, DRR, FJS, GM, GTC, HL, JE, JMZ, JR, MA, MAD, MCS, MM, NDO, PC, PP, RJH, SK, TM, and WJR performed analysis. AC, AMP, CSC, DRR, FJS, JMZ, MAD, MCS, and MWH supervised analysis. AC, AMW, DRR, GM, JMZ, PP, RH, SK, and WJR wrote the manuscript. All authors reviewed and approved the final manuscript.

## Acknowledgements

We would like to thank John R. Harting for assistance with HLA typing and Kristin A. Robertshaw for figure generation. SK and AMP were supported by the Intramural Research Program of the National Human Genome Research Institute, National Institutes of Health. This work utilized the computational resources of the NIH HPC Biowulf cluster (https://hpc.nih.gov).

This work was supported by NIH grant 1R01HG010040 to HL and NSFC grants 31571353 and 31822029 to JR. MCS is funded by the National Science Foundation (DBI-1350041) and National Institutes of Health (R01-HG006677). FJS and MM are funded by NIH grant UM1 HG008898.

This work utilized computational resources of DNAnexus and Google to apply DeepVariant to CCS reads.

Certain commercial equipment, instruments, or materials are identified to specify adequate experimental conditions or reported results. Such identification does not imply recommendation or endorsement by the National Institute of Standards, nor does it imply that the equipment, instruments, or materials identified are necessarily the best available for the purpose.

## Competing Financial Interests

AMW, AT, DRR, GTC, MWH, PP, RJH, WJR, and YQ are employees and shareholders of Pacific Biosciences. AC, AK, MAD, and PC are employees and shareholders of Google. AF and CSC are employees and shareholders of DNAnexus. AC is a shareholder and was an employee of DNAnexus for a portion of this work.

