## Supplementary Material for "Highly-accurate long-read sequencing improves variant detection and assembly of a human genome"

#### Detailed Author Contributions

CCS Library Preparation and Sequencing: DRR, PP, YQ

Quality Evaluation of CCS Reads: AMW, GM, RJH

Increased Mappability of CCS Reads: RJH

Small Variant Detection in CCS Reads: AC, AK, CSC, FJS, JMZ, MAD, NDO, PC, WJR

Phasing Small Variants: JE, TM, WJR

Improving Small Variant Detection with Haplotype Phasing: AC, AK, MAD, PC, WJR

Structural Variant Detection in CCS Reads: AMW, AT, FJS, HL, MCS, MA, MM

De Novo Assembly of CCS Reads: AF, AMP, AMW, DRR, JR, GTC, SK

Coverage Requirements for Variant Calling and De Novo Assembly: AC, AK, AMW, GTC, JE, TM, WJR

Revising and Expanding Genome in a Bottle Benchmarks: AMW, JMZ, NDO

### Supplementary Figures

#### Supplementary Figure 1

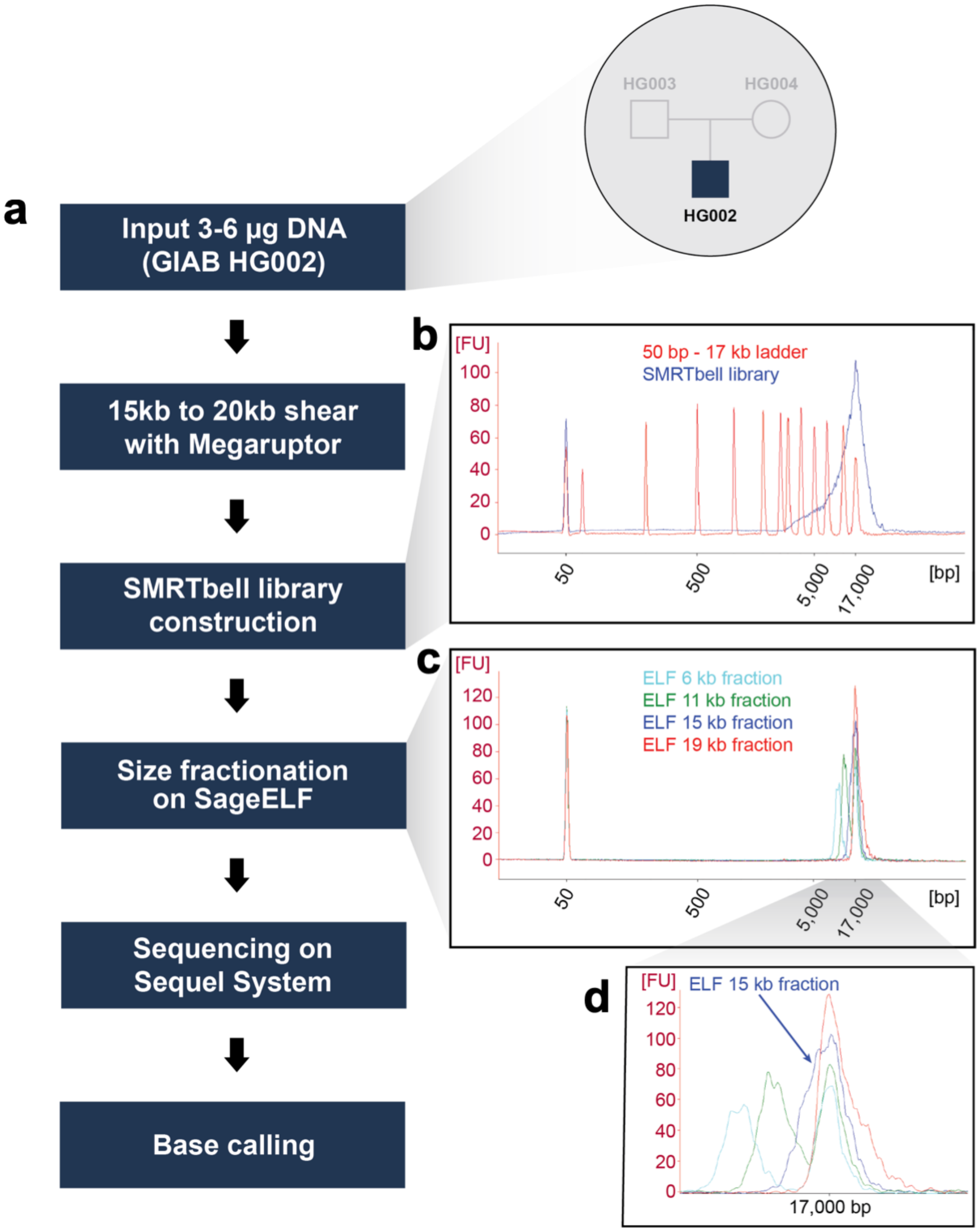


**Supplementary Figure 1. CCS protocol.** (a) Sample preparation and sequencing workflow. (b) BioAnalyzer trace for the SMRTbell library, sheared to target 15-20 kb fragments. "FU" is fluorescence units. (c) BioAnalyzer trace for ELF fractions of the SMRTbell library. (d) The fraction centered around 15 kb was used for sequencing.

#### Supplementary Figure 2

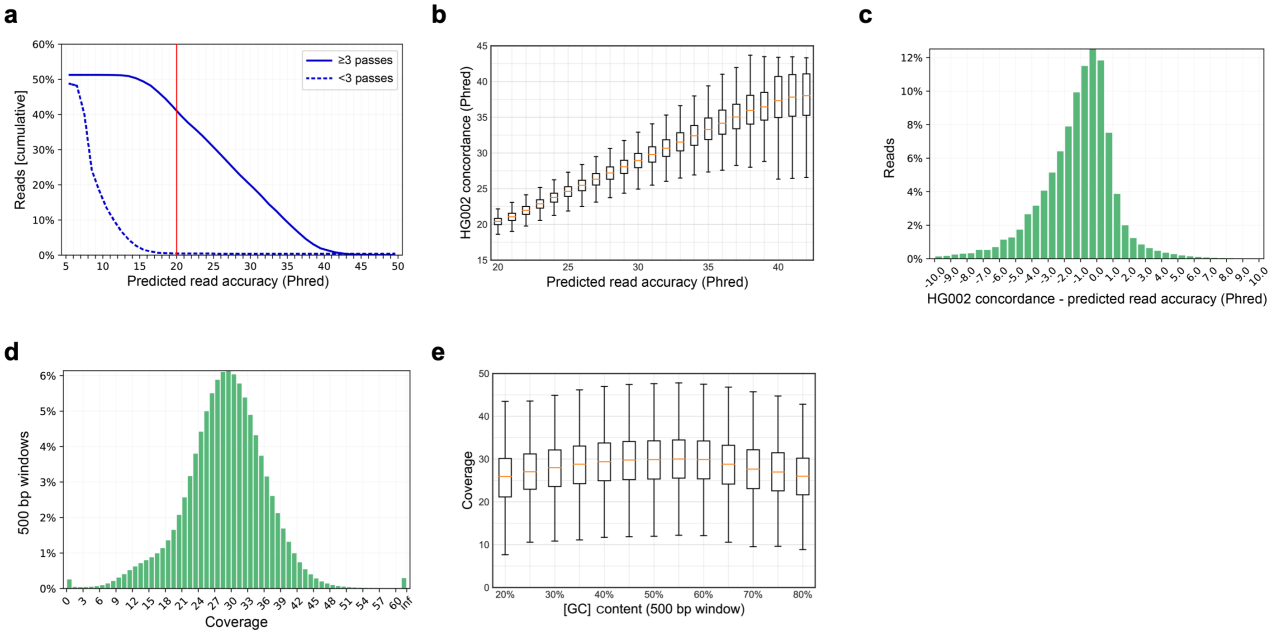


**Supplementary Figure 2. CCS read accuracy and coverage uniformity.**
(a) Distribution of accuracy predicted by the CCS algorithm for reads with fewer than 3 passes and at least 3 passes, which we consider a minimum pass count for CCS. Approximately half of reads have 3 or more passes; among those nearly all achieve Q20 predicted accuracy. (b) Distributions of HG002 concordance, measured against the GIAB benchmark, at levels of predicted read accuracy ($R^2$ of median = 0.9980), and (c) difference between concordance and predicted read accuracy show that the prediction is well-calibrated to the empirical concordance. (d) Distribution of coverage in 500 bp windows at non-gap positions in GRCh37. (e) Coverage distributions at levels of [GC] content, measured in 500 bp windows.

#### Supplementary Figure 3

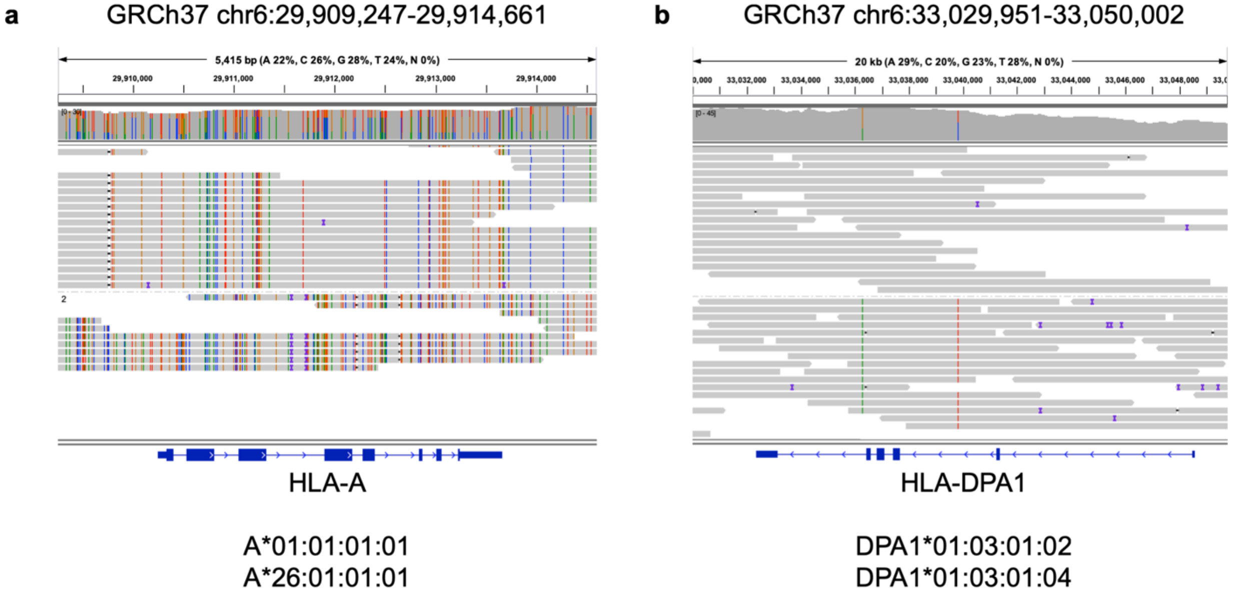


**Supplementary Figure 3. CCS read pileups at HLA genes.** The 13.5 kb CCS reads provide phasing and full four-field resolution of HLA class I and II genes[1], including (a) *HLA-A* for which HG002 has alleles that differ in the first field, and (b) *HLA-DPA1* for which HG002 has alleles that differ only in the fourth field from two intronic single nucleotide polymorphisms across 20 kb.

#### Supplementary Figure 4

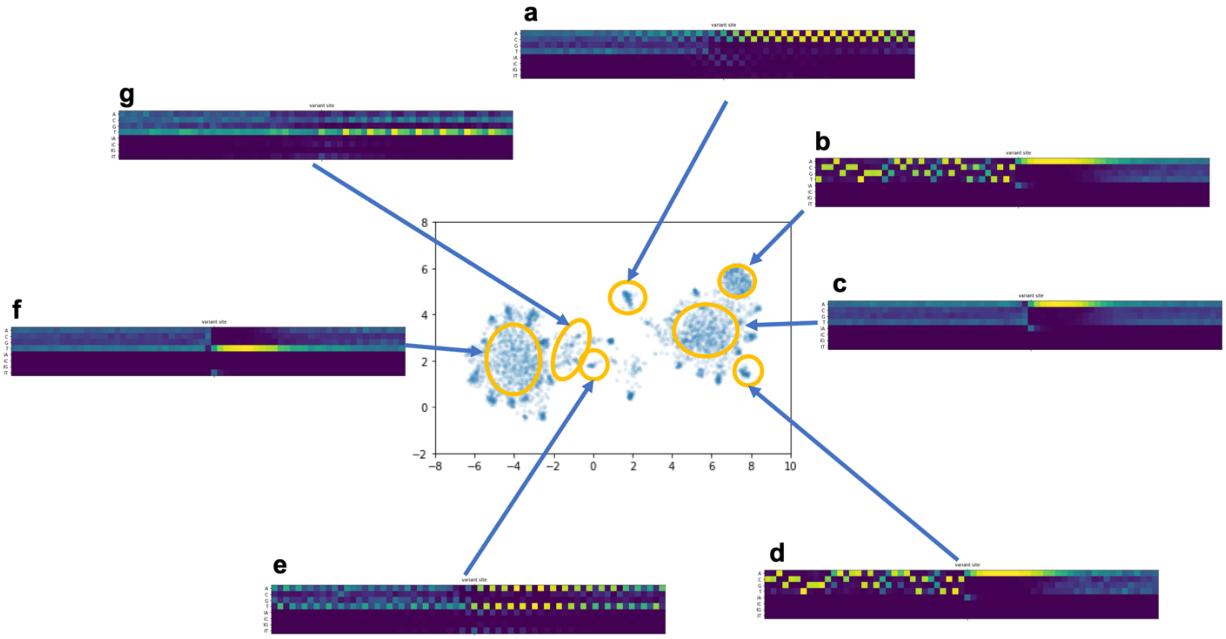


**Supplementary Figure 4. Alignment contexts around DeepVariant (CCS) discordances.** For all positions where the DeepVariant (CCS) callset is discordant with the GIAB benchmark, CCS read alignments to the position +/- 32 bp were encoded as a matrix (4 rows for A/C/G/T match, 4 rows for A/C/G/T insertion). The matrices were deflated into vectors with length 65×8=520 and embedded into two dimensions using UMAP[2]. Some distinct clusters represent simple, identifiable patterns: (a)(e) di-nucleotide repeats; (b)(d) poly-A tails of ALU elements[3]; (c)(f) homopolymer A/T runs without a specific prefix; and (g) [CT]-rich simple repeats.

#### Supplementary Figure 5

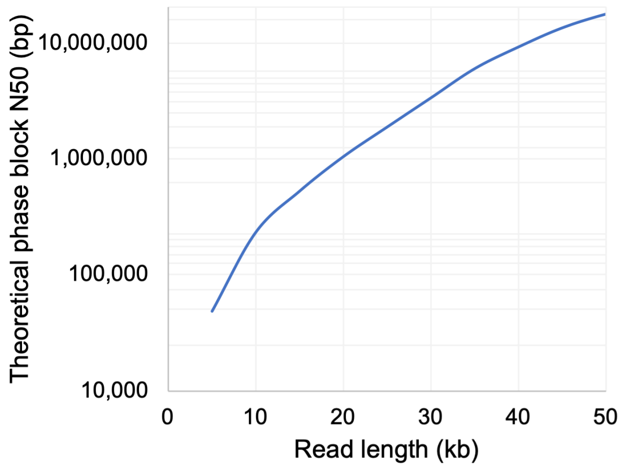


**Supplementary Figure 5. Theoretical phase block N50 in HG002 at different read lengths.** To model the phase blocks achievable with a given read length, cuts were introduced between heterozygous variants in the GIAB trio-phased HG002 variant callset that are separated by more than the read length, which effectively assumes that adjacent heterozygous variants separated by less than the read length can be phased.

#### Supplementary Figure 6

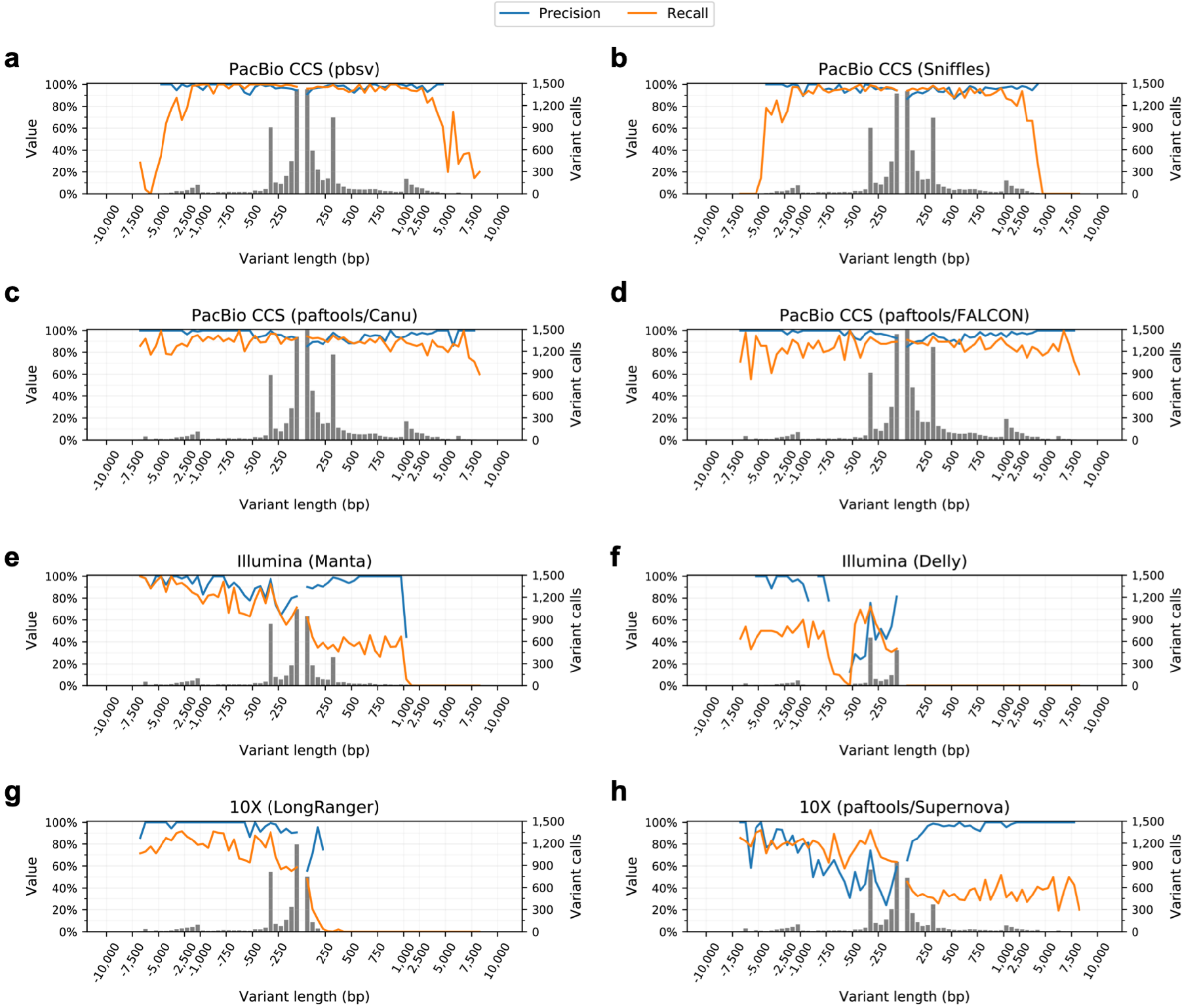


**Supplementary Figure 6. Structural variant calling performance.** Precision, recall, and number of variant calls in the GIAB benchmark regions for the PacBio CCS mapping-based variant callers (a) pbsv and (b) Sniffles; the PacBio CCS assembly-based callers (c) paftools/Canu (polished) and (d) paftools/FALCON (unpolished); the Illumina short-read callers (e) Manta and (f) Delly; and the 10X Genomics callers (g) LongRanger and (h) paftools/Supernova. Negative length indicates a deletion; positive length indicates an insertion. The histogram bin size is 50 bp for variants shorter than 1 kb, and 500 bp for variants >1 kb. Precision and recall are measured with Truvari against the GIAB benchmark.

#### Supplementary Figure 7

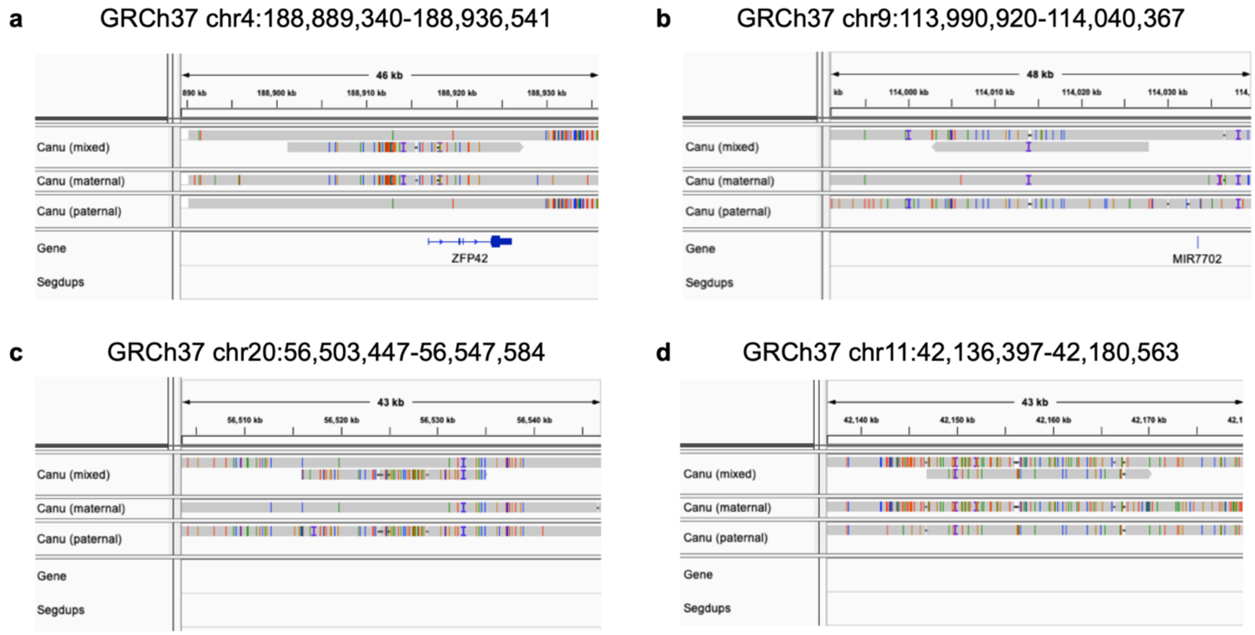


**Supplementary Figure 7. Haplotype resolution in the Canu mixed assembly.** The Canu mixed assembly is larger than the haploid human genome size because it resolves some heterozygous loci into separate maternal and paternal haplotypes.
(a) (b) Loci where the long primary contig matches the paternal haplotype and a smaller contig matches the maternal haplotype. (c) (d) Similar loci where the long primary contig matches the maternal haplotype and a smaller contig matches the paternal haplotype.

#### Supplementary Figure 8

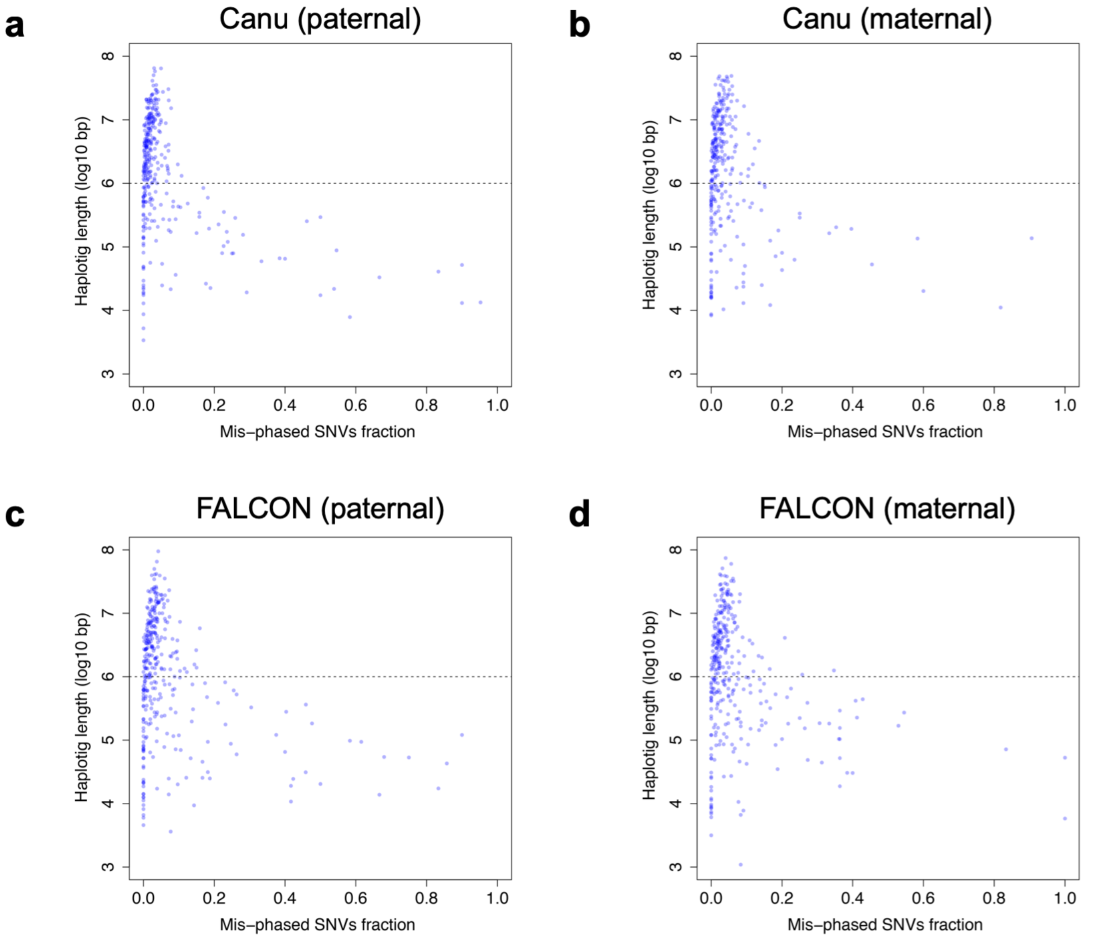


**Supplementary Figure 8. Mis-phasing analysis of parental assemblies**. Parent-specific heterozygous SNVs were identified in the GIAB benchmark callset. The "Mis-phased SNVs fraction" is the fraction of parent-specific SNVs from the wrong parent (e.g. $[SNV_{pat}]/[SNV_{pat}+SNV_{mat}]$ in a maternal contig). No large contigs have a high mis-phased SNVs ratio, which suggests proper phasing of the (a) Canu paternal, (b) Canu maternal, (c) FALCON paternal, and (d) FALCON maternal assemblies.

#### Supplementary Figure 9

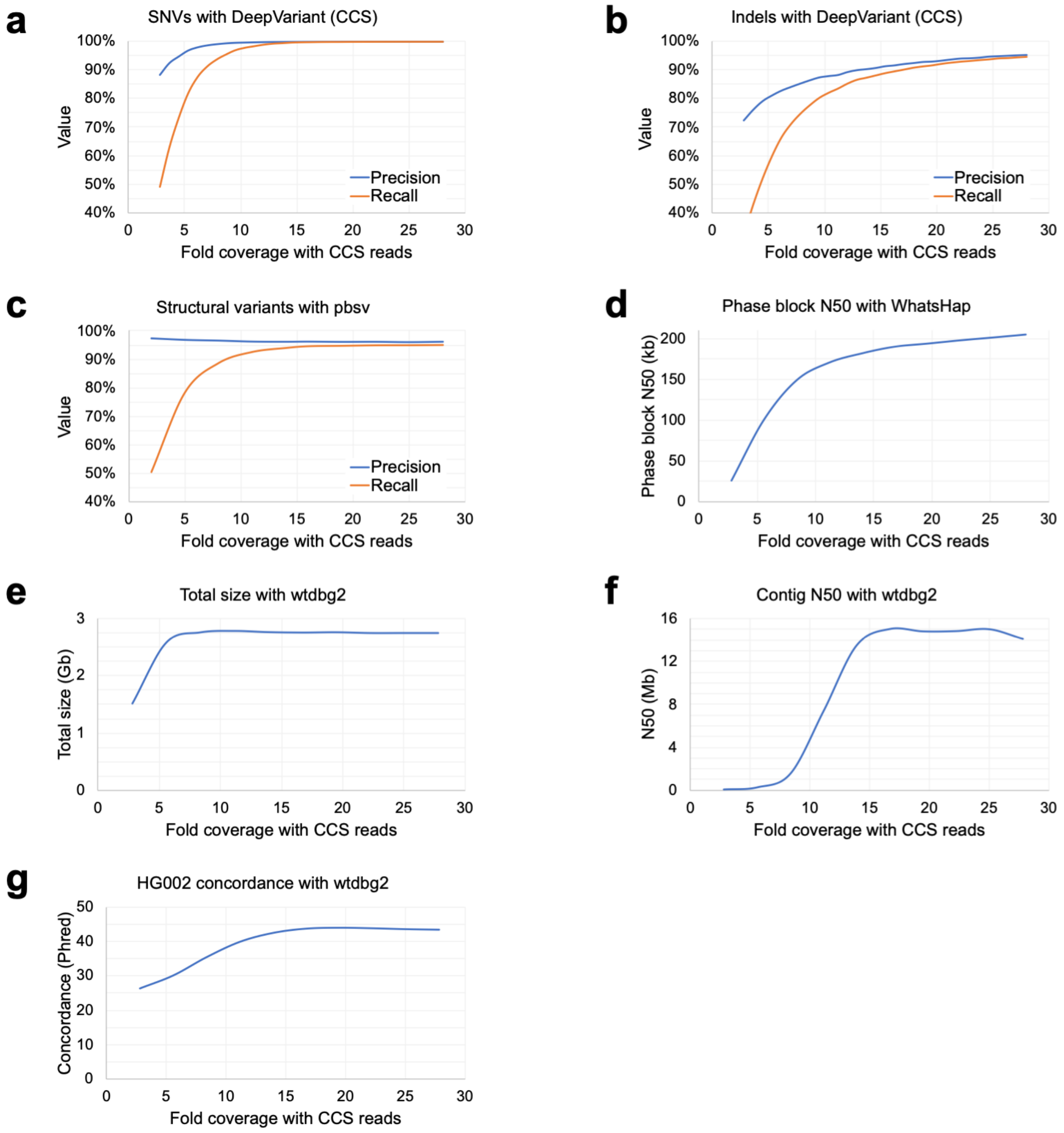


**Supplementary Figure 9. Coverage titration for variant calling, phasing, and assembly.** Precision and recall for (a) SNVs and (b) indels called with DeepVariant (CCS), subsampling in steps of 3%. (c) Precision and recall for structural variants called with pbsv, subsampling in steps of 10%. (d) Phase block N50 for phasing of the 28-fold DeepVariant (CCS) callset with WhatsHap, subsampling in steps of 10%. Phasing performance is similar with a callset produced at matched coverage (not shown). *De novo* assembly (e) completeness measured as total assembly size, (f) contiguity measured as contig N50, and (g) correctness measured as concordance to the HG002 GIAB benchmark for wtdbg2 assembly, subsampling reads in steps of 10%.

#### Supplementary Figure 10

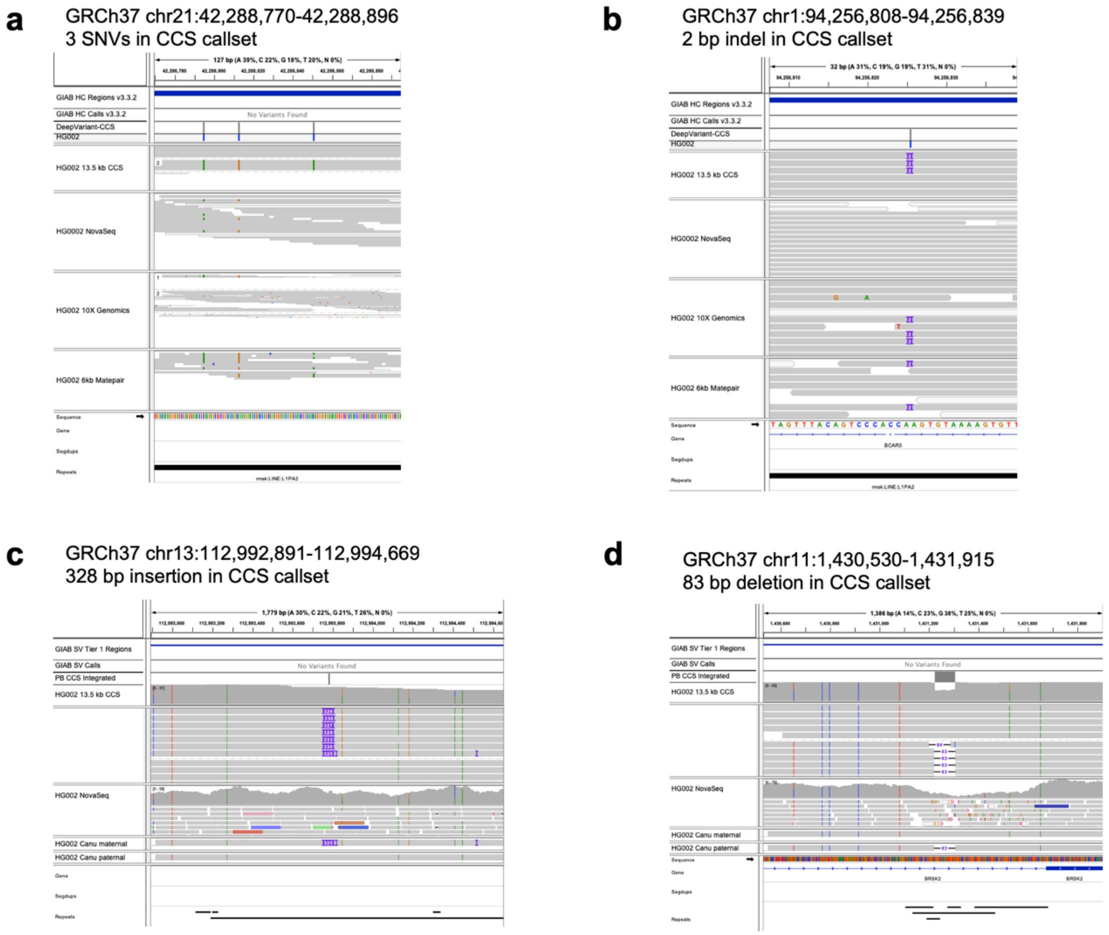


**Supplementary Figure 10. Likely errors in the GIAB benchmark identified by CCS callsets.** The high-quality GIAB benchmark and CCS variant callsets have strong, but not perfect, concordance. Manual curation of discrepancies identifies benchmark errors for all variant types that are correctable using the CCS variant callsets. Shown are four loci that the GIAB benchmark records as homozygous reference where CCS reads identify likely heterozygous variation: (a) Three SNVs supported by CCS reads and 6 kb matepair reads. (b) A 2 bp insertion supported by CCS reads, 10X Genomics reads, and 6 kb matepair reads. (c) A 328 bp insertion supported by CCS reads and assemblies. (d) An 83 bp insertion supported by CCS reads.

### Supplementary Tables

#### Supplementary Table 1

| | CCS reads | | | | NGS (NovaSeq) reads | | | |
|---|---|---|---|---|---|---|---|---|
| **Discordance** | **%** | **Freq. (1/bp)** | **Concordance** | | **%** | **Freq. (1/bp)** | **Concordance** | |
| Mismatch | 3.4% | 13,048 | 99.992% | (Q41) | 99.1% | 761 | 99.869% | (Q29) |
| Non-homopolymer indel | 4.6% | 9,669 | 99.990% | (Q39) | 0.3% | 241,876 | 99.999% | (Q54) |
| Non-homopolymer insertion | 3.6% | 12,359 | 99.991% | (Q41) | 0.1% | 884,457 | 99.999% | (Q59) |
| Non-homopolymer deletion | 1.0% | 44,425 | 99.998% | (Q46) | 0.2% | 332,920 | 99.999% | (Q55) |
| Homopolymer indel | 92.0% | 477 | 99.790% | (Q27) | 0.6% | 124,943 | 99.999% | (Q51) |
| Homopolymer insertion | 42.3% | 1,037 | 99.903% | (Q30) | 0.1% | 523,665 | 99.999% | (Q57) |
| Homopolymer deletion | 49.7% | 884 | 99.887% | (Q29) | 0.5% | 164,096 | 99.999% | (Q52) |
| **Total** | **100%** | **439** | **99.772%** | **(Q26)** | **100%** | **748** | **99.866%** | **(Q29)** |

**Supplementary Table 1. Discordances between read alignments and the HG002 GIAB benchmark for CCS and NGS reads.** An indel is considered a homopolymer event if the inserted/deleted basepairs match either the preceding or following reference basepair. “Percentage” is over all discordances, by type. "Freq." is frequency, or the number of read basepairs between discordances. “Concordance” considers only discordances of the given type. The "Q" value is concordance in Phred scale.

#### Supplementary Table 2

| Platform | Variant caller (training model) | SNVs Precision | SNVs Recall | SNVs F1 ^ | Indels Precision | Indels Recall | Indels F1 |
|---|---|---|---|---|---|---|---|
| Illumina (NovaSeq) | DeepVariant (Illumina model) | **<u>99.925%</u>** | **<u>99.940%</u>** | **<u>99.933%</u>** | **<u>99.450%</u>** | **<u>99.233%</u>** | **<u>99.341%</u>** |
| Illumina (NovaSeq) | GATK HaplotypeCaller (no filter) | 99.824% | 99.920% | 99.872% | 99.230% | 98.898% | 99.064% |
| PacBio (CCS) | DeepVariant (haplotype-sorted CCS model) | 99.778% | <u>99.937%</u> | 99.858% | 96.860% | 96.035% | 96.446% |
| PacBio (CCS) | DeepVariant (CCS model) | 99.807% | 99.904% | 99.855% | 95.387% | 94.501% | 94.942% |
| PacBio (CCS) | DeepVariant (Illumina model) | 99.533% | 99.793% | 99.663% | 23.991% | 81.692% | 37.090% |
| PacBio (CCS) | GATK HaplotypeCaller (hard filter) | 99.408% | 99.531% | 99.469% | 77.137% | 79.941% | 78.514% |

**Supplementary Table 2. Performance of small variant calling with CCS reads on chromosome 20.** DeepVariant models were not presented with chromosome 20 data before variant calling, so accuracy evaluations between GATK and DeepVariant are most comparable for chromosome 20. **Bold** indicates the highest value in each column. <u>Underline</u> indicates a value higher than the GATK HaplotypeCaller run on 30-fold Illumina NovaSeq reads. Coverage is 28-fold for PacBio CCS and 30-fold for Illumina NovaSeq. Callers are sorted ("^") based on F1 for SNVs.

#### Supplementary Table 3

| GRCh37 chrom | Heterozygous variants | % phased | Phase blocks | Phase block N50 (bp) | Hamming error rate | Switch errors | Switch error rate |
|---|---|---|---|---|---|---|---|
| 1 | 220,180 | 99.61% | 1,585 | 225,534 | 1.53% | 1,168 | 0.65% |
| 2 | 212,809 | 99.62% | 1,879 | 179,190 | 1.53% | 373 | 0.21% |
| 3 | 193,762 | 99.73% | 1,312 | 259,761 | 1.63% | 408 | 0.25% |
| 4 | 199,451 | 99.70% | 1,338 | 238,088 | 1.65% | 547 | 0.33% |
| 5 | 186,023 | 99.75% | 1,115 | 277,697 | 1.06% | 237 | 0.15% |
| 6 | 177,458 | 99.71% | 1,160 | 265,656 | 0.96% | 303 | 0.20% |
| 7 | 166,051 | 99.70% | 1,048 | 246,748 | 2.23% | 1,105 | 0.80% |
| 8 | 153,941 | 99.71% | 1,002 | 250,705 | 1.22% | 322 | 0.25% |
| 9 | 119,897 | 99.72% | 778 | 207,951 | 1.30% | 362 | 0.36% |
| 10 | 141,433 | 99.72% | 840 | 255,026 | 2.13% | 344 | 0.29% |
| 11 | 128,503 | 99.67% | 948 | 203,073 | 1.24% | 169 | 0.16% |
| 12 | 135,470 | 99.72% | 832 | 292,306 | 3.51% | 229 | 0.20% |
| 13 | 100,628 | 99.69% | 638 | 244,289 | 2.19% | 123 | 0.14% |
| 14 | 93,645 | 99.68% | 548 | 292,617 | 2.70% | 520 | 0.66% |
| 15 | 81,981 | 99.61% | 609 | 188,168 | 0.71% | 411 | 0.61% |
| 16 | 87,697 | 99.71% | 596 | 198,059 | 4.69% | 455 | 0.63% |
| 17 | 78,865 | 99.65% | 569 | 209,363 | 3.06% | 380 | 0.61% |
| 18 | 74,575 | 99.68% | 568 | 215,577 | 2.44% | 95 | 0.15% |
| 19 | 70,975 | 99.78% | 345 | 283,264 | 2.17% | 149 | 0.26% |
| 20 | 61,413 | 99.65% | 425 | 207,556 | 3.53% | 165 | 0.33% |
| 21 | 44,142 | 99.49% | 257 | 178,353 | 4.29% | 545 | 1.60% |
| 22 | 38,604 | 99.71% | 249 | 221,143 | 1.29% | 87 | 0.28% |
| **Autosomes** | **2,779,801** | **99.64%** | **19,215** | **206,063** | **1.91%** | **8,497** | **0.37%** |

**Supplementary Table 3. WhatsHap phasing performance on DeepVariant (CCS) callset.** WhatsHap provides highly complete phasing (99.64%) of heterozygous variants in the DeepVariant (CCS) callset that is concordant with the GIAB Trio/10X Genomics phasing benchmark set. Statistics are reported by WhatsHap with Hamming and switch error rates evaluated against the benchmark.

#### Supplementary Table 4

| | | All variants | | | Deletions | | | Insertions | | |
|---|---|---|---|---|---|---|---|---|---|---|
| **Platform** | **Caller** | *Prec.* | *Recall* | *F1 ^* | *Prec.* | *Recall* | *F1* | *Prec.* | *Recall* | *F1* |
| PacBio (CCS) | Integrated | 96.13% | **95.99%** | **96.06%** | **97.66%** | **96.88%** | **97.27%** | 94.97% | **95.30%** | 95.13% |
| PacBio (CCS) | pbsv | **96.26%** | 94.93% | 95.59% | 96.71% | 94.98% | 95.84% | **95.95%** | 94.89% | **95.42%** |
| PacBio (CCS) | Sniffles | 94.28% | 91.76% | 93.01% | 96.56% | 92.19% | 94.32% | 92.59% | 91.44% | 92.01% |
| PacBio (CCS) | paftools/Canu †‡ | 93.16% | 92.32% | 92.74% | 95.84% | 92.76% | 94.28% | 91.48% | 91.99% | 91.73% |
| PacBio (CCS) | paftools/FALCON † | 93.25% | 89.14% | 91.15% | 95.99% | 89.00% | 92.36% | 91.64% | 89.25% | 90.43% |
| Illumina | Manta | 85.34% | 55.88% | 67.53% | 85.95% | 76.90% | 81.17% | 92.12% | 39.65% | 55.44% |
| 10X | paftools/Supernova | 64.52% | 52.74% | 58.04% | 55.37% | 73.71% | 63.24% | 82.74% | 36.57% | 50.72% |
| 10X | LongRanger | 83.79% | 39.83% | 53.99% | 94.66% | 70.18% | 80.60% | 59.39% | 16.41% | 25.71% |
| Illumina | Delly | 65.92% | 19.90% | 30.58% | 65.92% | 45.70% | 53.98% | 0.00% | 0.00% | 0.00% |

**Supplementary Table 4. Structural variant calling performance.** Precision and recall are measured with Truvari against the GIAB benchmark. **Bold** indicates the highest value in each column; callers are sorted (“^”) based on F1 for all variants. † union of maternal and paternal assemblies; ‡ polished with Arrow.

#### Supplementary Table 5

| | | CPU core hours | | |
|---|---|---|---|---|
| **Haplotype** | **Assembler** | **Trio binning** | **Assembly** | **Arrow polishing** |
| Mixed | Canu | n/a | 2,136 | - |
| Mixed | FALCON | n/a | 2,650 | - |
| Mixed | wtdbg2 | n/a | 380 | - |
| Maternal | Canu | 350 | 751 | 71,226* |
| Maternal | FALCON | 350 | 1,683 | 26,137 |
| Maternal | wtdbg2 | 350 | 182 | - |
| Paternal | Canu | 350 | 841 | 70,069* |
| Paternal | FALCON | 350 | 1,568 | 26,183 |
| Paternal | wtdbg2 | 350 | 187 | - |

**Supplementary Table 5. CPU core hours for *de novo* assembly and polishing.** The CPU core hours required for trio binning, assembly, and polishing were recorded using the Unix `time` command. Assembly time includes read correction built into the assembler but excludes the total upfront CCS read generation (118,365 CPU core hours). The assemblers were run by different groups on different hardware, and thus times are not directly comparable. "*" Arrow polishing was run for one round on FALCON and two rounds on Canu; "n/a" = not applicable; "-" = not done

#### Supplementary Table 6

| | % reads assigned to haplotype | | |
|---|---|---|---|
| **k-mer (bp)** | **Maternal** | **Paternal** | **Unassigned** |
| 21 | 35.3% | 33.6% | 31.1% |
| 51 | 40.4% | 38.1% | 21.5% |
| 91 | 40.5% | 38.7% | 20.8% |

**Supplementary Table 6. CCS read classification by trio binning.** The percentage of CCS reads assigned to the maternal and paternal haplotype by k-mer size used in trio binning. CCS reads with an insufficient number of distinguishing k-mers are assigned to the “unassigned” haplotype, which includes reads from homozygous regions of genome.

#### Supplementary Table 7

| NCBI/ENA accession | Platform | Sample | Assembler + polish | Concordance ^ |
|---|---|---|---|---|
| - | PacBio (CCS) | HG002 (pat.) | Canu + Arrow | 99.9983% (Q47.7) |
| - | PacBio (CCS) | HG002 (mat.) | Canu + Arrow | 99.9981% (Q47.2) |
| - | PacBio (CCS) | HG002 | wtdbg2 | 99.9965% (Q44.6) |
| GCA_001542345 | PacBio (CLR) | HG002 | PBcR + Quiver | 99.9900% (Q40.0) |
| GCA_002077035 | PacBio (CLR) | HG001 | FALCON + Quiver | 99.9893% (Q39.7) |
| ERZ781176 | ONT + Illumina | HG001 | Canu + Nanopolish×2, Pilon×2, Racon×2 | 99.8694% (Q28.8) |
| - | ONT | HG001 | Canu + Nanopolish×2 | 99.6566% (Q24.6) |

**Supplementary Table 7. Reference concordance of assemblies from different platforms.** Concordance is measured against the GIAB HG002 benchmark. The three CCS read assemblies have higher concordance than accessioned assemblies provided with PacBio continuous long reads (CLR) or Oxford Nanopore read (ONT). ONT HG001 assembly is from https://obj.umiacs.umd.edu/marbl_publications/triobinning/albacore_canu_nanopolish2.fasta or https://bit.ly/2HBOjyq. "pat.": paternal, "mat.": maternal.

#### Supplementary Table 8

| | | Segdups | |
|---|---|---|---|
| **Haplotype** | **Assembler** | **Resolved (Mb)** | **Unresolved (Mb)** |
| Mixed | Canu | 63.6 | 111.8 |
| Mixed | FALCON | 46.1 | 129.3 |
| Mixed | wtdbg2 | 26.4 | 149.0 |
| Maternal | Canu | 60.2 | 115.2 |
| Maternal | FALCON | 43.2 | 132.2 |
| Maternal | wtdbg2 | 28.9 | 146.5 |
| Paternal | Canu | 60.0 | 115.4 |
| Paternal | FALCON | 41.7 | 133.7 |
| Paternal | wtdbg2 | 27.2 | 148.2 |

**Supplementary Table 8. Resolution of segmental duplications.** A segmental duplication in GRCh38 is considered resolved by an assembly if it is spanned by a contig with at least 50 kb on each flank, as measured by segDupPlots (https://github.com/mvollger/segDupPlots).

#### Supplementary Table 9

| Discrepancy | Variant type | Repeat family (if ≥1 kb) | Homopolymer length (bp) (if ≥6 bp) | Correct call | Chr | Position | Variant |
|---|---|---|---|---|---|---|---|
| AM | INDEL | | 19 | GIAB | 2 | 9,591,845 | CT/C |
| AM | INDEL | | | GIAB | 2 | 232,051,483 | GCA/GCATCATGGAGAATGGGACATCTC |
| AM | INDEL | | | GIAB | 3 | 37,083,407 | G/GA |
| AM | INDEL | | | CCS | 4 | 11,468,804 | CACACATATAT/C |
| AM | INDEL | L1PA2 | | CCS | 5 | 42,740,225 | CT/C |
| AM | INDEL | | Nearby 17 | GIAB | 6 | 41,984,320 | ACTAT/A |
| AM | INDEL | | 16 | GIAB | 8 | 73,675,279 | TAAAA/T |
| AM | INDEL | | 13 | GIAB | 13 | 76,646,445 | G/GA |
| AM | INDEL | | 16 | GIAB | 15 | 44,350,983 | C/CA |
| AM | INDEL | | 13 | GIAB | 19 | 1,586,670 | A/ATTT |
| AM | SNP | HERVH-int | | CCS | 2 | 5,143,996 | G/A |
| AM | SNP | | | GIAB | 2 | 230,174,543 | A/G |
| AM | SNP | L1PA2 | | CCS | 4 | 165,276,021 | T/C |
| AM | SNP | | 8 | GIAB | 5 | 16,287,108 | A/C |
| AM | SNP | | 8 | GIAB | 11 | 41,384,344 | C/T |
| AM | SNP | | 20 | GIAB | 12 | 51,793,781 | A/C |
| AM | SNP | | 9 | GIAB | 13 | 34,840,815 | G/T |
| AM | SNP | L1PA3 | | CCS | 13 | 48,291,499 | A/C |
| AM | SNP | | 12 | GIAB | 13 | 71,512,745 | A/T |
| AM | SNP | | | GIAB | 21 | 25,668,597 | G/A |
| FN | INDEL | | | GIAB | 1 | 162,491,859 | A/ATGTCTAG |
| FN | INDEL | | 12 | GIAB | 2 | 152,262,374 | G/GTT |
| FN | INDEL | | 14 | GIAB | 2 | 236,062,930 | G/GTT |
| FN | INDEL | L1PA2 | 9 | GIAB | 3 | 107,982,543 | AT/A |
| FN | INDEL | | 18 | GIAB | 4 | 149,672,221 | A/ATT |
| FN | INDEL | | | CCS | 8 | 5,930,728 | TACAC/T |
| FN | INDEL | | 6 | GIAB | 10 | 29,087,199 | T/TCC |
| FN | INDEL | | | GIAB | 15 | 26,120,981 | C/CTTACACTGGGCTTTTTGTAAGGA |
| FN | INDEL | | | CCS | 15 | 41,943,823 | T/TCCTCTTCTCTCCTCTCC |
| FN | INDEL | | 15 | GIAB | 17 | 5,198,683 | C/CA |
| FN | SNP | | 16 | GIAB | 5 | 55,201,041 | A/G |
| FN | SNP | | | GIAB | 6 | 8,353,625 | C/T |
| FN | SNP | | | CCS | 6 | 9,737,425 | T/C |
| FN | SNP | | | GIAB | 6 | 57,283,620 | T/C |
| FN | SNP | | 13 | GIAB | 7 | 135,981,582 | T/A |
| FN | SNP | | | CCS | 7 | 157,385,671 | A/G |
| FN | SNP | | | GIAB | 9 | 117,917,190 | A/C |
| FN | SNP | | 13 | GIAB | 9 | 129,471,234 | T/A |
| FN | SNP | | 5 | CCS | 17 | 32,064,214 | A/G |
| FN | SNP | | 25 | GIAB | 17 | 68,021,050 | T/A |
| FP | INDEL | L1PA2 | | CCS | 1 | 94,256,825 | A/AAC |
| FP | INDEL | L1HS | | CCS | 2 | 153,864,971 | AT/A |
| FP | INDEL | L1M2 | 13 | GIAB | 3 | 97,014,398 | AT/A |
| FP | INDEL | L1HS | 7 | CCS | 4 | 112,819,087 | GA/G |
| FP | INDEL | L1PA2 | | CCS | 4 | 165,026,074 | A/AG |
| FP | INDEL | | 10 | GIAB | 6 | 64,897,720 | A/AT |
| FP | INDEL | | 15 | GIAB | 7 | 38,338,238 | C/CA |
| FP | INDEL | | | GIAB | 8 | 132,575,025 | C/CAAAAAAAAA |
| FP | INDEL | L1P1 | | CCS | 11 | 23,338,682 | C/CT |
| FP | INDEL | | 20 | GIAB | 11 | 61,993,476 | CA/C |
| FP | SNP | L1HS | | CCS | 1 | 35,034,071 | T/C |
| FP | SNP | L1HS | | CCS | 3 | 79,181,734 | C/T |
| FP | SNP | | Nearby 8 | GIAB | 4 | 55,520,593 | G/A |
| FP | SNP | L1HS | 7 | CCS | 4 | 94,532,444 | T/G |
| FP | SNP | ALR/Alpha | | CCS | 8 | 46,873,565 | C/T |
| FP | SNP | | 11 | GIAB | 9 | 6,900,971 | C/T |
| FP | SNP | L1PA2 | | CCS | 9 | 22,350,168 | A/C |
| FP | SNP | | 13 | GIAB | 20 | 1,347,896 | A/G |
| FP | SNP | | 12 | GIAB | 20 | 4,159,335 | C/T |
| FP | SNP | L1PA2 | | CCS | 21 | 42,288,851 | C/A |

**Supplementary Table 9. Manual curation of small variant discrepancies between CCS callset and GIAB benchmark.** For the "Discrepancy" column, "AM" means genotype difference, "FN" means false negative (in benchmark but not callset), and "FP" means false positive (in callset but not benchmark). "Repeat family" column is from the RepeatMasker track from the UCSC Genome Browser. "Correct call" column is "GIAB" when the benchmark was deemed correct by expert curators, and "CCS" when the CCS callset was deemed correct. Rows where the correct call is from the CCS callset are colored blue.

#### Supplementary Table 10

| Discrepancy | Variant type | Length (bp) | Simple repeat length (bp) (if ≥100 bp) | Simple repeat period (bp) | Correct call | Chr | Position |
|---|---|---|---|---|---|---|---|
| FN | DEL | -32,196 | | | GIAB | 1 | 152,555,543 |
| FN | DEL | -2,269 | | | GIAB | 2 | 159,958,799 |
| FN | DEL | -49,058 | 172 | 71 | - | 4 | 34,779,881 |
| FN | DEL | -127 | 466 | 127 | GIAB | 4 | 123,733,539 |
| FN | DEL | -357 | 1,921 | 20 | GIAB | 13 | 30,131,788 |
| FN | DEL | -108 | 589 | 54 | GIAB | 13 | 114,841,327 |
| FN | DEL | -52 | | | GIAB | 16 | 85,800,468 |
| FN | DEL | -565 | 1,403 | 561 | - | 19 | 4,884,873 |
| FN | DEL | -55 | | | GIAB | 19 | 57,683,315 |
| FN | DEL | -120 | 917 | 40 | GIAB | 20 | 62,510,913 |
| FN | INS | 62 | | | GIAB | 2 | 228,113,946 |
| FN | INS | 104 | 163 | 27 | GIAB | 3 | 66,992,107 |
| FN | INS | 52 | | | GIAB | 3 | 172,678,665 |
| FN | INS | 125 | 230 | 23 | GIAB | 5 | 105,107,607 |
| FN | INS | 727 | 651 | 65 | - | 6 | 40,459,830 |
| FN | INS | 172 | 815 | 4 | - | 9 | 135,394,538 |
| FN | INS | 6,179 | | | GIAB | 12 | 71,053,961 |
| FN | INS | 51 | 125 | 3 | GIAB | 13 | 29,161,602 |
| FN | INS | 3,268 | | | GIAB | 14 | 67,862,850 |
| FN | INS | 58 | 472 | 33 | CCS | 19 | 14,488,489 |
| FP | DEL | -1,432 | | | - | 1 | 108,735,819 |
| FP | DEL | -50 | | | CCS | 2 | 65939,406 |
| FP | DEL | -80 | 1,274 | 16 | CCS | 6 | 167,162,349 |
| FP | DEL | -80 | 332 | 40 | CCS | 7 | 129,149 |
| FP | DEL | -65 | 632 | 168 | - | 10 | 134,253,963 |
| FP | DEL | -83 | 333 | 83 | CCS | 11 | 1,431,223 |
| FP | DEL | -74 | 1,674 | 74 | CCS | 12 | 6,038,958 |
| FP | DEL | -128 | | | - | 13 | 107,435,844 |
| FP | DEL | -63 | 1,227 | 22 | GIAB | 17 | 230,498 |
| FP | DEL | -300 | 1,923 | 60 | GIAB | 18 | 77,569,248 |
| FP | INS | 103 | | | CCS | 4 | 141,283,453 |
| FP | INS | 52 | 588 | 26 | CCS | 4 | 190,329,327 |
| FP | INS | 202 | 893 | 18 | - | 8 | 146,172,196 |
| FP | INS | 176 | 1,184 | 24 | - | 10 | 132,840,681 |
| FP | INS | 783 | 1,184 | 24 | - | 10 | 132,841,387 |
| FP | INS | 328 | | | CCS | 13 | 112,993,782 |
| FP | INS | 60 | 312 | 4 | CCS | 16 | 85,867,748 |
| FP | INS | 54 | 527 | 18 | - | 17 | 10,662,861 |
| FP | INS | 84 | | | CCS | 18 | 53,029,667 |
| FP | INS | 267 | 641 | 37 | CCS | X | 67,035,046 |

**Supplementary Table 10. Manual curation of structural variant discrepancies between CCS callset and GIAB benchmark.** For the “Discrepancy” column, “FN” means false negative (in benchmark but not callset), and “FP” means false positive (in callset but not benchmark). “Simple repeat length” and “Simple repeat period” are from the simpleRepeat track from the UCSC Genome Browser. “Correct call” column is “GIAB” when the benchmark was deemed correct by expert curators, “CCS” when the CCS callset was deemed correct, and “-“ when it is unclear which callset is correct (typically due to complex tandem repeats that permit multiple representations of the same variant). Rows where the correct call is from the CCS callset are colored blue.
